# Direct testing of natural twister ribozymes from over a thousand organisms reveals a broad tolerance for structural imperfections

**DOI:** 10.1101/2024.07.11.603121

**Authors:** Lauren N. McKinley, McCauley O. Meyer, Aswathy Sebastian, Benjamin K. Chang, Kyle J. Messina, Istvan Albert, Philip C. Bevilacqua

## Abstract

Twister ribozymes are an extensively studied class of nucleolytic RNAs. Thousands of natural twisters have been proposed using sequence homology and structural descriptors. Yet, most of these candidates have not been validated experimentally. To address this gap, we developed CHiTA (Cleavage High-Throughput Assay), a high-throughput pipeline utilizing massively parallel oligonucleotide synthesis and next-generation sequencing to test putative ribozymes *en masse* in a scarless fashion. As proof of principle, we applied CHiTA to a small set of known active and mutant ribozymes. We then used CHiTA to test two large sets of naturally occurring twister ribozymes: over 1, 600 previously reported putative twisters and ∼1, 000 new candidate twisters. The new candidates were identified computationally in ∼1, 000 organisms, representing a massive increase in the number of ribozyme-harboring organisms. Approximately 94% of the twisters we tested were active and cleaved site-specifically. Analysis of their structural features revealed that many substitutions and helical imperfections can be tolerated. We repeated our computational search with structural descriptors updated from this analysis, whereupon we identified and confirmed the first intrinsically active twister ribozyme in mammals. CHiTA broadly expands the number of active twister ribozymes found in nature and provides a powerful method for functional analyses of other RNAs.

**GRAPHICAL ABSTRACT:** 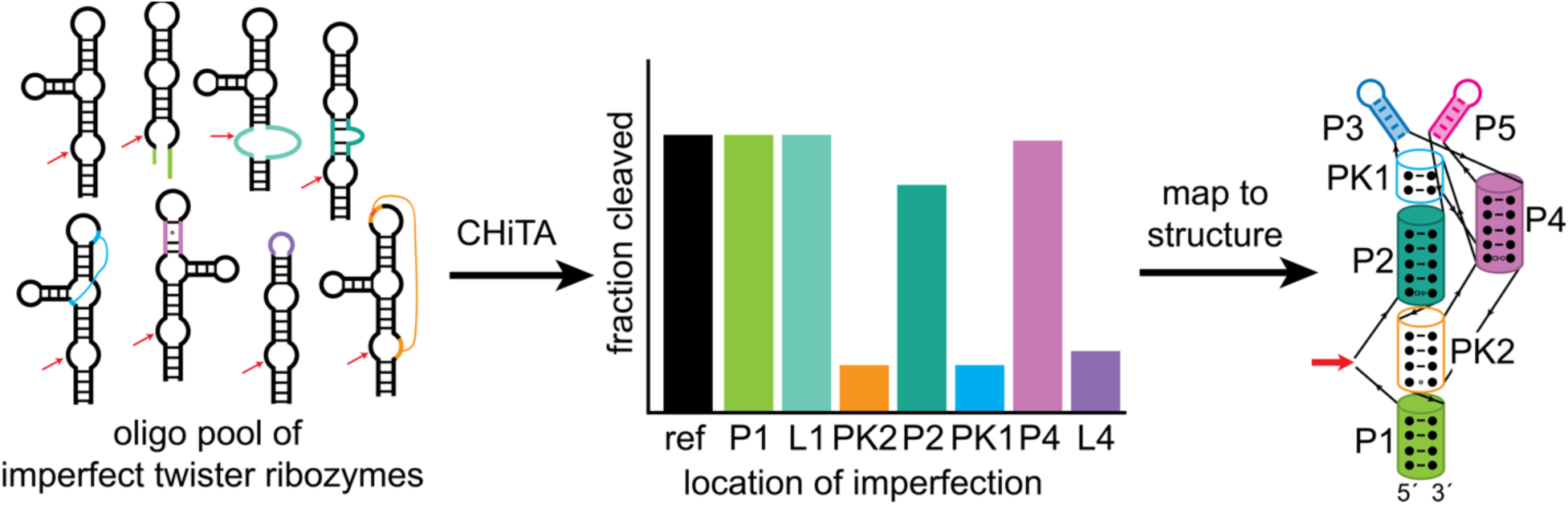

## INTRODUCTION

For many classes of RNA, their function is derived from their structure. Unlike DNA, which assumes a simple double-stranded helix, RNA adopts a diverse array of intramolecular structural features including hairpins, internal loops, and multihelical junctions, which aid in the functionality of the RNA (1–3). Interestingly, RNA can tolerate imperfections such as bulges, deletions, insertions, mismatches, mutations, and overhangs while maintaining its function (4). This structural diversity allows RNA to fulfill biological roles apart from being a conduit for genetic information (2, 3). Riboswitches and RNA thermometers, for instance, employ conformational switching to regulate transcription and translation in response to small molecules and temperature, respectively (5, 6). RNA enzymes, known as ribozymes, catalyze protein synthesis and splicing as well as use cleavage reactions to mediate gene expression and RNA half-life (7).

Broadly, RNA enzymes can be classified by size as either small (50-150 nt) or large (>150 nt) (7, 8). Small nucleolytic ribozymes are a type of catalytic RNA that self-cleave their phosphodiester backbone via a general acid-base mediated transphosphorylation reaction, which yields a 5′- and a 3′-cleavage fragment with a terminal 2′, 3′-cyclic phosphate and 5′-hydroxyl group, respectively (7, 9, 10). At least 10 classes of small nucleolytic ribozymes have been described: hammerhead (11), hairpin (12), Hepatitis Delta Virus (HDV) (13), Varkud satellite (Vs) (14), *glmS* (15), HDV-like (16), twister (17), twister sister (18), pistol (18), and hatchet (18). While many of the structural and mechanistic properties of these ribozyme classes have been elucidated (19), their biology is not deeply understood. They are ubiquitously distributed throughout all three domains of life, as thousands of candidates have been predicted (7).

Programs to identify putative ribozymes have been developed as a result of advances in bioinformatics (9). For example, the Basic Local Alignment Search Tool (*BLAST*) (20) relies upon sequence homology, while programs like *RNAMotif* (21), *Infernal* (22), and *RNArobo* (23) employ descriptions of the secondary structure to search for different ribozyme classes. In applying these programs, Roth et al. described 2, 690 putative twister ribozymes (17), and Lee et al. more recently detected 10, 181 ribozymes from 9 classes in 2023 (24). *RfamGen* (25) was developed to design ribozymes by assimilating both sequence similarity and secondary structure with machine learning, whereby it generated 1, 000 sequences belonging to the *glmS* family (25). Despite these advances, very few putative ribozymes have been evaluated experimentally Several (HT) methodologies have been developed that are capable of testing thousands of mutant ribozymes *en masse* (26). Using doped solid phase synthesis or error-prone PCR to produce the DNA templates and next-generation sequencing (NGS) to measure cleavage, these pipelines examined all single and double mutants of one ribozyme from a given class (26–32). Adaptations, like *k*-seq, allow for kinetic rate profiling with high accuracy if timepoints are taken (30). Together these methods have yielded systematic and comprehensive means for assaying the influence of mutations on catalysis (26). While powerful, these methods typically test only one ribozyme, and thus do not consider the full structural diversity observed in many classes of ribozymes found in nature, including circular permutations (17, 33–38), auxiliary stem-loops (17, 36), bulges, deletions, insertions, and overhangs.

To address this gap, massively parallel oligonucleotide synthesis (MPOS) can be employed to create a pool containing thousands of ribozymes (39–42). Oligo pools are created as single-strand DNA (ssDNA) typically on the scale of picomoles of each sequence, necessitating PCR amplification and transcription to produce RNA (39–42). Consequently, non-native PCR primer binding sites and a transcriptional promoter are added to the template DNA. These pools have been utilized in screenings for RNA modifications (41), optimal regulatory elements (enhancers, promoters, etc.) (43, 44), RNA structural studies (45), and ribozyme self-cleavage assays focused on designing and engineering ribozymes (25, 46). One challenge posed by the oligo pool approach, though, is the addition of non-native flanking sequences, which if transcribed, may result in alternative conformations that disrupt canonical folding of the RNA. In the case of small self-cleaving ribozymes, such flanking sequence has been reported to alter ribozyme behavior (47–49) and may preclude accurate assessment of the intrinsic reactivity of the ribozyme. Indeed, for ribozyme assays that employ MPOS (25, 46), non-native flanking sequences were not removed. Removal of non-native sequence in a scarless fashion is therefore desirable.

Herein, we report the development of a Cleavage High-Throughput Assay (CHiTA), which quantifies activity for thousands of natural ribozyme candidates *en masse*, in a scarless fashion, using MPOS and NGS. On a small-scale to establish the method, we observed the expected extent of cleavage for 8 ribozymes from 3 classes and saw that cleavage was specific to known cleavage sites. CHiTA was then implemented on a large-scale with 1, 613 putative twister ribozymes previously predicted in literature (17) as well as 1, 012 novel candidate twisters that we predicted in over 1, 000 organisms. Here, we focus on type-P1 circular permutes of twister ribozymes, but we consider all three type of auxiliary stem-loops (P0/L0, P3/L3, and P5/L5). Comprehensive analysis of the length and imperfections (number, location, and type) within each structural element of twister ribozymes demonstrated a surprising tolerance for imperfections in helical and loop elements. We then constructed descriptors informed by our structural analysis, which led us to identify the first intrinsically active twister in a mammal.

## MATERIALS AND METHODS

### Data analysis, availability, and reproducibility

Putative ribozymes were predicted using the BLASTn webserver (https://blast.ncbi.nlm.nih.gov/Blast.cgi) (20), Rfam (50), and the *R2DT* webserver (https://rnacentral.org/r2dt) (51). Command line tools utilized included the following: *bwa mem* (v. 07.17) (52), *Entrez Direct* (v. 19.7) (53), NCBI’s *Command Line Tools* (v. 16.4.5) (53), the *Infernal* package (v. 1.1.4) (22), R2R (v. 1.0.6) (54), *RNArobo* (v. 2.1.0) (23), the *RNAStructure* package (v. 6.3) (55), *SAMtools* package (v. 1.15) (56). Python (v. 3.6) (57) was employed to filter and count reads from sequencing data. The R coding language (v. 4.3.2) (58) and RStudio interface (v. 2023.06.0+421) (59) were used to reformat text files, perform statistical analysis, and create graphics. Additional R dependences employed included: *cowplot* (v. 1.1.1), *dplyr* (v. 1.1.2), *extrafont* (v. 0.19), *ggcorrplot* (v. 0.1.4.1), *ggdist* (v. 3.3.0), *ggplot2* (v. 3.4.2), *ggpubr* (v. 0.6.0), *ggrepel* (v. 0.9.3), *grid* (v. 4.3.2), *patchwork* (v. 1.1.3), *png* (v. 0.1-8), *stringr* (v. 1.5.0), *tidyverse* (v. 2.0.0), *R2easyR* (v. 0.1.1) (https://github.com/JPSieg/R2easyR), *rstatix* (v. 0.7.2), *stats* (v. 4.3.2), and *viridis* (v. 0.6.3). For the Computational Discovery Pipeline, the *SnakeMake* (60) management program was employed, and genomes were downloaded from RefSeq Database (61). Code was made with assistance from *GitHub Copilot* (v. 1.180.0) and *ChatGPT* (v. 3.5).

### Statistical analysis

Linear regression was performed in R with the *ggcorrplot* package (v. 0.1.4.1) to fit data to the line of best fit. Calculations for the goodness of fit (R^2^) and Pearson’s correlation coefficient (r) were completed in R with the *stats* package (v. 4.3.2). Statistical significance was conducted using the *rstatix* (v. 0.7.2) package in R. For comparison of just two groups, the Mann-Whitney U test was employed. For comparison of three or more groups, the Kruskal-Wallis test, was applied, followed by a Games-Howell post hoc test to correct for bias encountered in performing multiple tests. The Mann-Whitney U and Kruskal-Wallis nonparametric tests were chosen to account for the non-uniform distribution of data and unequal sample size (62). All p-values are reported with 2 significant figures and are provided in respective SI tables. Only p-values with n ≥10 are shown in the figures.

### Reagents

Materials used in this study are as follows. All oligo pools, DNA templates, and primers were from Integrated DNA Technologies (IDT). The Phusion High Fidelity (HF) Buffer (B0518S), Phusion DNA Polymerase (M0530S), rCutSmart Buffer (B6004S), Bsal-HF v2 (R3733S), 6X DNA Loading Dye (B7024S), DNase I (M0303S), DNA Adenylation Kit (B2610S), Mth RNA Ligase (M2611A), ATP for adenylation (N0757A), PEG8000 (B10045), 1X T4 RNA ligase buffer (B0216S), and T4 RNA ligase “2 Truncated” (M0242S) were from New England Biolabs. The 100 mM dithiothreitol (DTT) (707265ML) and SuperScript III (12574026) used for reverse transcription were from Thermo Fisher Scientific. The Nucleospin PCR clean-up kit (740609) and NTC buffer (740654) were by Macherey-Nagel. The Circligase II kit (CL9021K) was from Lucigen; the RNA Clean and Concentrator kit (R1019) was from Zymo Research; and SYBR Gold (S11494) was from Invitrogen. The [α-^32^P] ATP (0.250 mCi, 500000017275) was from Revvity. The T7 polymerase was prepared in house.

### Computational discovery of new twister candidates for the novel dataset

Twister ribozyme candidates for the novel oligo pool were found by searching the 16 nt consensus sequences of P4 and L4, 5′-CCGGTCCCAAGCCCGG-3′ (17), using the BLASTn webserver (Figure S1) (20). Organisms entered into BLASTn were limited to those with genomes in the Nucleotide Collection and Whole-genome Shotgun Contig databases found in BLAST, and all other parameters were default. Sequences with < 10 nt of sequence identity were discarded. Next, the 5′ and 3′ surrounding sequences were manually inspected for structural elements consistent with twister ribozymes including pseudoknots (“NC” for PK1, “GRG” for PK2_3’_), predicted cleavage site (“NA”), and catalytic residues (“A” general acid and “G” general base) (17, 63, 64). To assess whether these sequences were novel, they were checked against published twister ribozyme sequences from Roth et al. (17), Liu et al. (65), Lee et al. (24), Eckert et al. (38), and Rfam (twister-P1, RF03160) (50). Novel sequences were then folded with default parameters on the *R2DT* webserver (51), which can identify a known functional RNA and then orient the secondary structure on the page in a standard format. A majority of sequences conformed to the twister ribozyme template within *R2DT*. Those that did not were folded either by *Fold* from the RNAStructure package (v. 6.3) (55) with default settings or by visual inspection. Usually, sequences folded with *Fold* adhered to the canonical, non-pseudoknotted secondary structure of twister ribozymes, with the exception of P1. All sequences and genetic contexts are reported in Supplementary File 1.

### CHiTA Experimental Characterization

#### Preparation of oligo pool containing ribozyme candidates

Ribozymes were ordered as ssDNA in oligo pools from IDT. Each ribozyme had forward and reverse PCR primer binding sites with a T7 promoter. To facilitate transcription, 1 or 2 non-native Gs were added immediately upstream of ribozymes whose sequence began with 1 or 0 Gs, respectively. Included within the reverse primer binding site was a BsaI restriction digest site that allowed for removal of the reverse primer binding site and the BsaI site itself, providing a “scarless” end for transcription (Figure 1, Figure S2).

**Figure 1.**
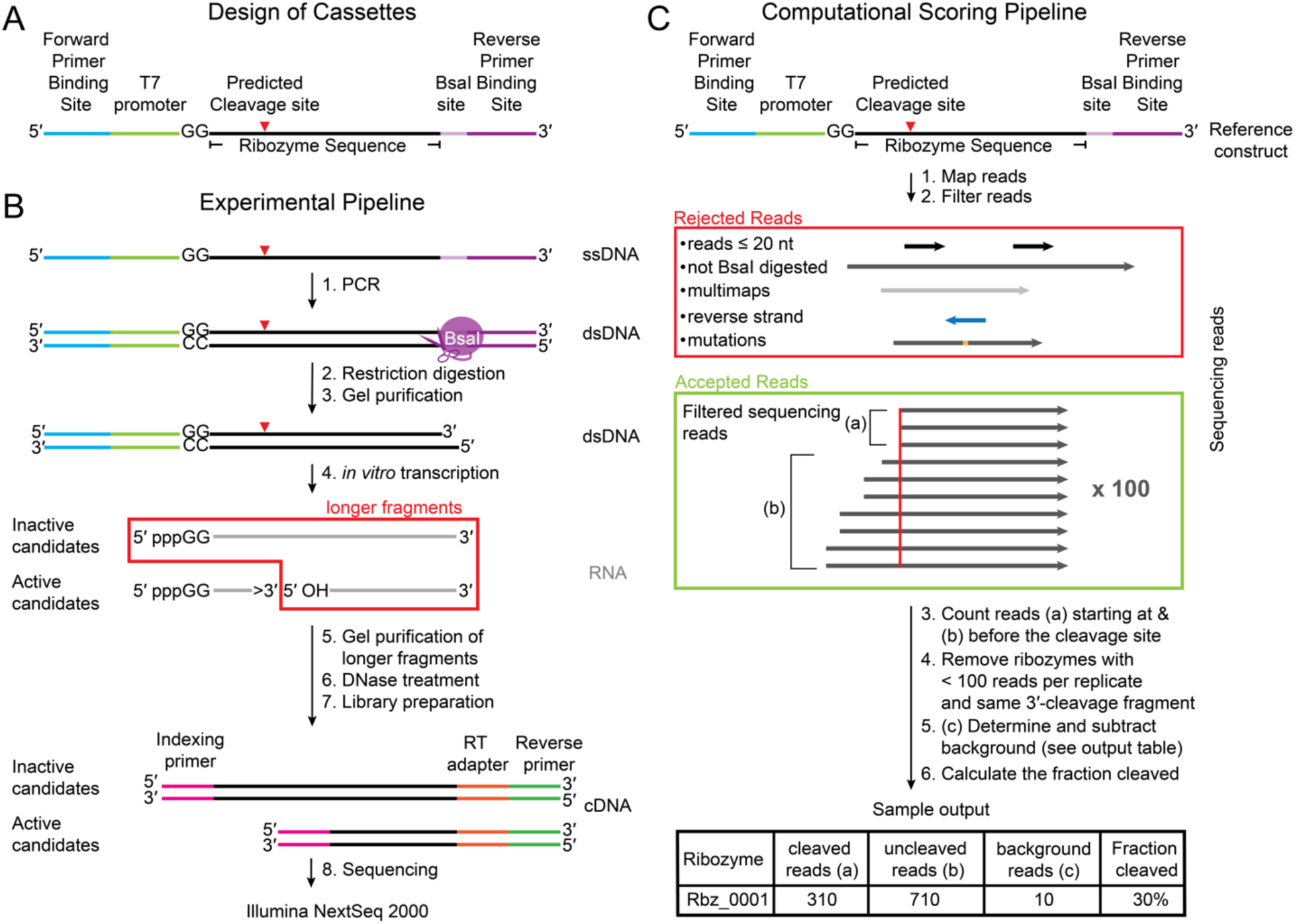
Overview of the CHiTA pipeline. CHiTA was used for testing the activity of small self-cleaving ribozymes *en masse*. (A) Design of cassettes into which putative ribozymes were inserted. Cassettes included common forward and reverse PCR primer binding sites, a T7 promoter, and a BsaI restriction site. B) Experimental Pipeline starting from an oligo pool that underwent PCR amplification, a restriction digest, *in vitro* transcription, and library preparation. (C) Computational Scoring Pipeline in which reads were filtered, counted, corrected for background, and used to determine the fraction cleaved for each ribozyme. (B-C) The red arrow denotes the predicted cleavage site of the ribozyme.

In total, 3 oligo pools were ordered: The small-scale oligo pool containing 8 ribozymes (50 pmol of each template DNA), and the two large-scale oligo pools containing the 1, 613 candidate twister ribozymes from literature (17) and the 1, 012 novel candidate twister ribozymes that we discovered (10 pmol of each template DNA). Sequences for all oligo pools are provided in Supplementary File 2.

#### PCR amplification of template ssDNA

ssDNA sequences from the oligo pools were PCR amplified into double stranded DNA (dsDNA) with Phusion HF DNA Polymerase under the following conditions: 0.2 mM each dNTP, 0.5 µM each forward and reverse PCR primers, 1 ng of template ssDNA, 1X Phusion HF buffer, and 2 U of Phusion polymerase (New England Biolabs). Reactions were initially denatured at 98 °C for 30 s, then denatured at 98 °C for 10 s, annealed at 62 °C for 1 min, and extended at 72 °C for 20 s for 25 three-temperature cycles before a final extension at 72 °C for 5 min. To achieve three technical replicates, PCR reactions were conducted in triplicate for each pool and samples were never combined. Each of the three PCR reactions was column purified with a Nucleospin PCR clean-up kit according to the manufacturer’s protocol (Macherey-Nagel). The concentration of dsDNA from each PCR reaction was determined on a microvolume UV-vis spectrometer (DeNovix).

#### Bsal restriction digest to remove the reverse PCR primer site

Once the DNA concentration was determined, the dsDNA was restriction digested with BsaI to completely remove the reverse PCR primer bindng site. The restriction digestion was done on each of the technical replicates under the following conditions: 1X rCutSmart buffer, 3-5 µg of dsDNA from PCR, and 20 U of BsaI-HF v2 (New England Biolabs), incubated at 37 °C for 16 h to drive the reaction. Reactions were mixed with 6X DNA loading dye (New England Biolabs) to a final 1X concentration. We note that even after optimization, the yield of BsaI restriction reactions was only ∼75%. As such, digested dsDNA was gel purified by running an 8% non-denaturing 1X TBE polyacrylamide gel electrophoresis (PAGE) for 4 h. The gel was stained with SYBR Gold (Invitrogen), and bands were visualized under UV light. Product bands were excised, and the corresponding gel slices were crushed and soaked overnight in TEN_250_ (10 mM Tris pH 7.5, 1 mM EDTA, 250 mM NaCl) at 4 °C, and then ethanol precipitated. After ethanol precipitation, the dsDNA concentration was measured on a microvolume UV-vis spectrometer (DeNovix).

#### *In vitro* transcription of digested template DNA

Digested template DNA was *in vitro* transcribed in triplicate with recombinant T7 RNA polymerase in the following conditions: 40 mM Tris (pH 8.2), 2 mM dithiothreitol (DTT), 1 mM spermidine, 4 mM ATP, 4 mM CTP, 4 mM UTP, 8 mM GTP, 25 mM MgCl_2_, 1-2 µg of digested template DNA, and 6% w/v T7 polymerase (prepared in house). All NTP solutions were adjusted to pH 8.2 using 1 M Tris prior to transcription. An elevated concentration of GTP was used to improve the efficiency of transcription initiation (66). Reactions were incubated at 37 °C for 3-3.5 h, with additional 6% w/v T7 polymerase added at 1.5 h to aid RNA production. Aliquots were removed at 1.5 and 3.5 h for the small-scale oligo pool only. For the large-scale oligo pools, transcription was terminated at 3 h. Unlike *glmS* ribozymes (29), twister ribozymes cleave co-transcriptionally and so active ribozymes would have cleaved during *in vitro* transcription (17). Reactions were quenched by adding an equal volume of 2X formamide loading buffer (95% formamide, 0.1X TBE, 20 mM EDTA, 0.025% bromophenol blue). Full-length ribozymes and the larger 3′-cleavage fragments were separated from the much smaller 5′-cleavage fragments using PAGE with a 10% denaturing gel (8.3 M urea) run at 40 W for 3 h. The gel was stained with SYBR Gold (Invitrogen), and bands were visualized under UV light. The longer fragments were excised from the gel and crushed and soaked overnight in TEN_250_ at 4°C, before being ethanol precipitated the next day. Subsequently, the concentration of the collected RNA was measured on a microvolume UV-Vis spectrometer (DeNovix). To remove DNA, template RNA was loaded onto an RNA Clean and Concentrator column (Zymo Research) and treated with DNaseI (New England Biolabs) according to the manufacturer’s instructions. Purified RNA was eluted from the column and quantitated on a microvolume UV-Vis spectrometer (DeNovix).

#### CHiTA Library Preparation

Library preparation generally followed previously reported protocols from our lab (67–69). See the Supplementary Materials and Methods for details.

### CHiTA Computational Characterization

#### Mapping to the reference genome

FASTQ files containing raw sequencing data were first mapped to their respective reference genome using *bwa mem* (v. 07.17) (52). The resulting SAM files were then converted to BAM files using *samtools sort* (v. 1.15) and indexed using *samtools index* (v. 1.15) (56).

#### Filtering of reads

Reads were filtered and counted using a custom python script (Supplementary File 3). Settings were adjusted to discard reads containing primer and promoter sequences, although two nucleotides of the reverse primer were allowed to account for non-templated 3′-end addition by the T7 polymerase (70). The following reads were filtered out: 1. short reads (≤ 20 nt), 2. secondary and supplementary reads, 3. reads that multi-mapped (mapping quality = 0), 4. reads that aligned to the reverse strand, and 5. reads with any difference from the reference sequence.

#### Counting reads

By definition, self-cleavage of ribozymes occurs between positions -1 and +1. Then, we counted all reads that passed filtering, including cleaved reads (reads that start at position 1, “*R*_clv_”), and uncleaved reads (all reads that start upstream of position 1, “*R*_unclv_”) for each ribozyme. To establish background and specificity of cleavage, reads starting at positions -3, -2, -1, 2, 3, and 4 were counted and compared to reads starting at position 1.

Ribozymes with fewer than 100 combined cleaved and uncleaved reads in each replicate were not analyzed to improve accuracy (30). All values are reported in Supplementary File 4.

#### Background correction

We used reads flanking the cleavage site to account for background. First, for each library the percentage of background reads occurring at position i (*P*_i_) upstream (i=-3, -2, or -1) or downstream (i=2, 3, or 4) of the cleavage site was determined according to equation 1:

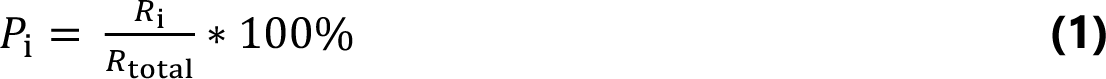

where *R*_i_ was the number of filtered reads starting at the i^th^ position, and *R*_total_ was all filtered reads. *P*_i_ was averaged across all ribozymes and then across positions i = -3, -2, -1, 2, 3, and 4 to determine the percentage of background (*P*_bkgd_) (Table S1). The number of background reads (*R*_bkgd_) for each ribozyme was calculated according to equation 2:

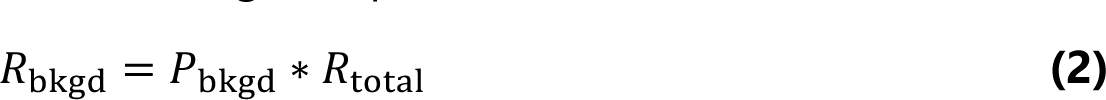

where *P*_bkgd_ was multiplied by *R*_total_ for each ribozyme. Background-corrected uncleaved reads (*R*_unclv, corr_) for each ribozyme were determined according to equation 3:

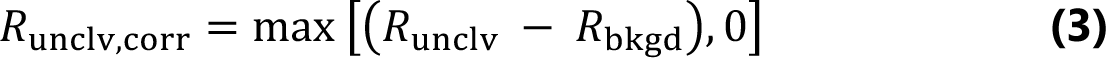

where *R*_bkgd_ for a ribozyme was subtracted from the raw number of all uncleaved reads (*R*_unclv_) for that ribozyme. Likewise, the background-corrected cleaved reads (*R*_clv, corr_) for each ribozyme were determined according to equation 4:

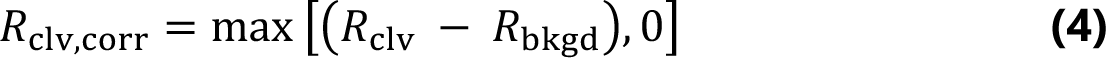

where *R*_bkgd_ for a ribozyme was subtracted from the raw number of cleaved reads (*R*_clv_) for that ribozyme. Both equations 3 and 4 set the minimal number of reads at 0. This method for correcting background was used over subtracting a single number of reads from all ribozymes due to variable coverage of the ribozymes. All values are reported in Supplementary File 4.

#### Calculating fraction cleaved

Using the background-corrected number of cleaved and uncleaved reads, the fraction cleaved (*f*_cleaved_) was calculated for each ribozyme according to equation 5:

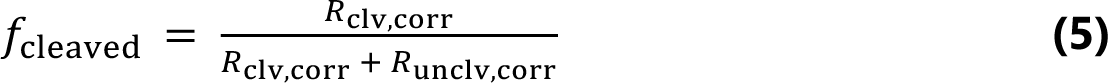

where background-corrected cleaved reads (*R*_clv, corr_) were divided by the total number of background-corrected counted reads (*R*_clv, corr_ plus *R*_unclv, corr_). *f*_cleaved_ was then averaged across the three replicates for each ribozyme. All values are reported in Supplementary File 4.

#### Partitioning of reads

To determine if cleavage was specific to the cleavage site, the number of background-corrected reads (*R*_i, corr_) was first calculated for positions adjacent to the cleavage site according to equation 6:

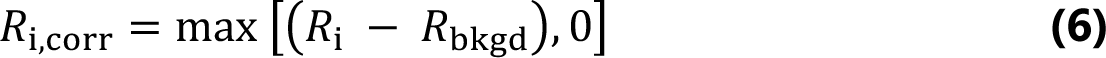

where *R*_bkgd_ for a ribozyme was subtracted from the raw number of reads at position i (*R*_i_). *R*_i, corr_ was calculated at positions -3, -2, -1, 2, 3, and 4. The background-corrected reads from positions -3, -2, -1, 1 (cleavage site), 2, 3, and 4 were then summed (Q) according to equation 7:

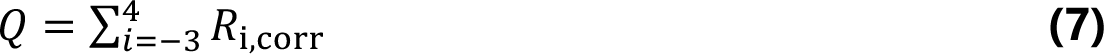

The fraction of reads at each site was finally calculated according to equation 8,

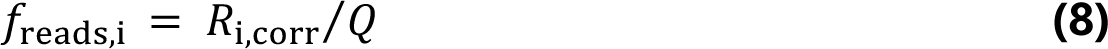

Significance was limited to 2 decimal places. All values are reported in Supplementary File 4.

#### Co-transcriptional cleavage assays

To verify that CHiTA provided an accurate measurement of *f*_cleaved_, 17 ribozymes with wide-ranging CHiTA-calculated *f*_cleaved_ values, were selected for conventional gel-based co-transcriptional cleavage assays. The T7 promoter was added upstream of each ribozyme along with 1 or 2 non-native Gs for ribozymes whose sequence began with 1 or 0 Gs, respectively, to facilitate transcription. For Dre 1-3 the sequence “5′-GGAGA-3′” was added in place of these non-native G’s. No additional downstream sequences (i.e reverse PCR primer binding site) were added. The reverse complement was then purchased from Integrated DNA Technologies (IDT) as ssDNA. All sequences tested are listed in Table S2 as their DNA template and corresponding RNA transcript. To prepare a hemi-duplex for transcription, 10 µM ssDNA template and 10 µM T7 promoter (5′-TAATACGACTCACTATAG-3′) in 25 mM NaCl were heat denatured at 90 °C for 2 min and then annealed at room temperature for ∼10 min. Each ribozyme was then transcribed in triplicate at 37 °C for 3 h as per above with the following exceptions: the ATP concentration was supplemented with 5 µCi [α-^32^P] ATP (Revvity, 0.250 mCi), and we used 1 µM hemi-duplex DNA template. Prior to transcription, all NTP solutions were adjusted to pH 8.2 using 1 M Tris. Timepoints (4 µL) were removed at 1, 5, 10, 15, 30, 60, 90, 120, and 180 min then quenched in 25 µL of loading buffer (95% formamide, 20 mM EDTA, 0.1X TBE, 0.1% w/v xylene cyanol, and 0.025% w/v bromophenol blue). From each timepoint, 5 µL was fractionated using PAGE with a 10% denaturing (8.3 M urea) gel run at 20 W for ∼2 h. Gels were then dried and imaged using a Phosphorimager (Typhoon GE Healthcare). Intensities from the full-length and 3′-cleavage fragment were quantified using ImageQuantTL (v. 8.2). The *f*_cleaved_ was calculated according to equation 9:

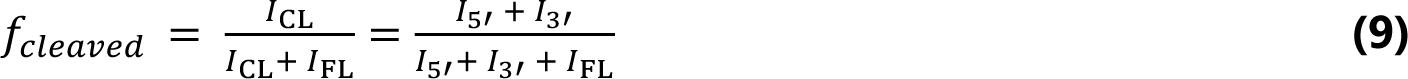

where *I*_CL_, *I*_5′_, *I*_3′_, and *I*_FL_ are the background-subtracted intensities of the cleaved, 5′-, and 3′-cleavage fragments, and full-length transcript, respectively. In most cases *I*_5′_ was ignored because the 5′-cleavage fragment was too short to detect or produced no signal because it had no As. Replicates were then averaged together for each ribozyme.

This same assay was used to test mammalian candidate ribozymes Hsap-1-1, Hsap-1-2, and Ttru-1-1. Template DNA was prepared as described above and the sequences are provided in Table S2 as both the DNA template and RNA transcript. Reactions occurred under the aforementioned conditions, with timepoints removed at 5, 10, 30, 60, 120, 180, 240, and 1440 min. The *f*_cleaved_ was calculated according to equation 9.

#### Structural analysis of twister ribozymes

Imperfections in the structural elements of putative ribozymes were assessed by first aligning all sequences to the secondary structure model for twister ribozymes. For twister ribozyme candidates predicted in literature, the alignment was downloaded as a stockholm file from Roth et al. (17). For new twister ribozyme candidates, sequences were aligned using *cmbuild* and *cmalign* from the *Infernal* package (v. 1.1.4) (22). First, references sequences for the twister-P1 family (RF03160) were downloaded from Rfam (50) as a stockholm file. This was then utilized in *cmbuild* (22) to construct a model to align candidate twisters. Default parameters were used. Putative ribozymes were aligned to the reference model using *cmalign* (22) where the -- nonbanded and --notrunc flags were utilized to generate the optimally accurate alignment and to specify that no sequences were truncated, respectively. Results were output as a non-interleaved stockholm file, and probabilities of covariation were hidden using the --noprob flag.

A custom R script was employed to determine the structural imperfections of all aligned twister candidates. Sequences pertaining to each structural element (P0/L0, linker, P1, L1, GA, GB, P2, PK1, L2, P3/L3, P4, L4, PK2, P5/L5) were first extracted. The length of each element was determined. For certain paired elements (P1, P2, P4, PK2), the predicted ΔG°_37_ was calculated using *BiFold* from the RNAStructure package (v. 6.3) (55) with default settings. Similarly, for auxiliary stem-loops (P0/L0, P3/L3, and P5/L5), the predicted ΔG°_37_ was determined for the minimum free energy (MFE) structure utilizing *Fold* from the RNAStructure package (v. 6.3) (55) with default settings. These values were imported into the script. Note that for PK1, no graphs depicting the length or predicted ΔG°_37_ are shown because PK1 was two nucleotides for all ribozymes tested, which was too short to calculate the predicted ΔG°_37_. Subsequently, the number, location, general and specific type of imperfections were noted. For paired elements, imperfections included bulges, deletions, insertions, mismatches, mutations, and overhangs. For loops, imperfections included deletions, insertions, and mutations. Due to the large variation observed in auxiliary stem-loops no imperfections were noted. After assessing individual elements, the whole ribozyme sequence was examined. The length and ΔG°_37_ of the entire ribozyme were obtained as described above. Combinations of imperfections, including the location, number, and general type of imperfections, were also denoted. Findings were then exported as a csv file and are reported in Supplementary File 5.

#### Sequence alignment of select twister ribozymes

Select twister ribozymes were aligned using the Clustal Omega webserver (https://www.ebi.ac.uk/jdispatcher/msa/clustalo) (v. 1.2.4) (71). Sequences were input in FASTA file format and default parameters were employed.

#### Determination of ribozyme consensus sequence and structure

To compute consensus sequences of twister ribozymes, we created four separate FASTA files containing active ribozymes (*f*_cleaved_ > 0.20): 1. ribozyme without a P3/L3 or P5/L5, 2. Ribozymes with a P3/L3, 3. Ribozyme with a P5/L5, and 4. all ribozymes. Each fasta file was aligned to the consensus model as described above using *Infernal* (v. 1.1.4) (22). The resulting stockholm files were input into the RScape webserver (http://eddylab.org/R-scape/) (55, 72–74). Default parameters were employed, and the “improve given structure” option was selected.

#### Computational pipeline to search for additional twister ribozymes

Informed by our structural analysis, we developed a Computational Discovery Pipeline (Figure S3) to search for additional ribozyme variants using *RNArobo* (23), *Infernal* (22), and *R2R* (54). Input files included structural descriptors of desired ribozyme variants, a list of genomes to search, and a reference stockholm file containing the structural alignment. Separate structural descriptors were designed in the format necessitated by *RNArobo* (23) and were based upon our structural analysis for twister ribozymes with no auxiliary stem-loops, a P3/L3, or a P5/L5 (Figure S4). Genomes of select organisms were downloaded from the RefSeq database (61) as FASTA files using *Entrez Direct* (v. 19.7) (53) and NCBI’s *Command Line Tools* (v. 16.4.5) (53). The reference Stockholm file was generated using *cmbuild* (22) as described above. Using a *SnakeMake* wrapper, the pipeline first employed *RNArobo* (v. 2.1.0) (23) to search input genomes for sequences matching the structural descriptors. Flags -c and -u were used to search both strands of the genome and avoid overlapping sequences. Next, sequences were aligned to the reference structural model for twister ribozymes using *cmalign* (22) with the same parameters as aforementioned. Finally, sequences were folded using *R2R* (54) with default parameters. Raw sequences were output in the form of a FASTA file, aligned sequences as a Stockholm file, and folded structures as a pdf. To minimize false positives, we only reported sequences with a bit score higher than 0.

## RESULTS

### Development of Cleavage High-Throughput Assay (CHiTA)

To evaluate thousands of putative natural ribozymes simultaneously, we developed a HT assay that we termed “CHiTA” for Cleavage High-Throughput Assay, which uses MPOS and NGS. Leveraging the large-scale opportunities afforded by MPOS, we devised cassettes into which candidate ribozymes could be inserted (Figure 1A, S2). These cassettes had common forward and reverse PCR primer binding sites and a T7 promoter. Within the 5′-end of the reverse primer binding site was a BsaI restriction site, which allowed the reverse primer binding site to be entirely removed prior to transcription, providing for a scarless transcript (Figure S2, row 2). Initial iterations of the pipeline included a permanent reverse primer, which had the opportunity to interact with the ribozyme and induce alternate conformations. Such structural changes could artificially alter ribozyme activity and prevent accurate assessment of the intrinsic reactivity of these ribozymes (48, 49). Our pipeline uniquely averted this issue by ensuring that no non-native flanking sequences were transcribed (46, 25). Candidate sequences were inserted into the cassettes and ordered as ssDNA in an oligo pools(see Materials and Methods).

In the Experimental Pipeline (Figure 1B), the oligo pool was first amplified by PCR to obtain sufficient material for *in vitro* transcription. A Bsal restriction digestion was then performed to remove the entire reverse PCR primer binding site and Bsal site (Figure S2, row 2). After which, the digested products were selected using gel purification (Figure 1B). Next, the dsDNA was *in vitro* transcribed, whereupon active ribozymes cleaved co-transcriptionally. After transcription, 3′-cleavage fragments and uncleaved full-length ribozymes, which are of similar length, were selected via denaturing PAGE (Figure 1B, red enclosure). To prepare the RNA for sequencing, we employed a similar library preparation to protocols previously reported from our lab (67–69). We utilized the shared 3′-terminal hydroxyl of the 3’-cleavage fragments and full-length ribozymes to ligate on an adenylated reverse transcription primer. The RNA was then reverse transcribed using SuperScript III. Following reverse transcription, the cDNA was circularized using CircLigase II and amplified using indexing PCR. Libraries were then sequenced with Illumina Next-Seq 2000 as 150 bp single-end reads.

In order to quantify the extent of ribozyme cleavage, we devised a Computational Scoring Pipeline to calculate *f*_cleaved_ for each ribozyme (Figure 1C). First, reads were aligned to the reference genome, which contained sequences of all putative ribozymes from the oligo pool. Next, the resulting BAM files were filtered using a custom python script to remove reads that were short (≤ 20 nt) and undigested, as well as those that multi-mapped, mapped to the reverse strand, or had any mutations. From the remaining reads, we counted the number of cleaved and uncleaved reads for each ribozyme. Cleaved reads were defined as those that started at the 5′-end of the 3′-cleavage fragment (nucleotide 1), while uncleaved reads were those that started at any nucleotide upstream of nucleotide 1 (Figure 1C). Ribozymes that produced the same 3′-cleavage fragment or whose replicates had < 100 reads were discarded (30). We then background-corrected the number of cleaved and uncleaved reads to account for enzyme stalling, random breakage of the RNA backbone, and other possible events (Equations 1-4, Table S1). The *f*_cleaved_ for each ribozyme was then calculated by dividing the number of background-corrected cleaved reads by the total number of background-corrected cleaved and uncleaved reads, then averaging across the replicates (Equation 5).

### Validation of CHiTA with ribozymes from several classes

We began by testing CHiTA on a small-scale group of 8 ribozymes from 3 different classes, using active and inactive variants. For the active ribozymes, we included twister ribozymes env-9 and env-22 (17), as well as a hammerhead ribozyme from *Schistosoma mansoni* (75) (Figure 2A-C), all of which have previously been shown to self-cleave *in vitro* (75–77). For the inactive twister mutants, we turned to mutational studies which have demonstrated that changes to catalytic bases (general acid, general base) are deleterious to self-cleavage efficacy (28, 63, 76–78). As such, we mutated both A1 (general acid) and G57/47 (general base) to Cs in env-9 (Figure 2A, magenta) and env-22 (Figure 2B, magenta). To inactivate the hammerhead ribozyme, we mutated an internal loop to form a continuous helix in Stem I (Figure 2C, magenta), which has been shown to prevent self-cleavage by disrupting tertiary interactions (75). Finally, as a negative control, we included the human HDV-like CPEB3 ribozyme with its 9 nt inhibitory upstream flanking sequence (Figure S5A) (48). From this, we also created an intrinsically inactive mutant, via a C57U mutation at the general acid (Figure S5A, magenta) (79).

**Figure 2.**
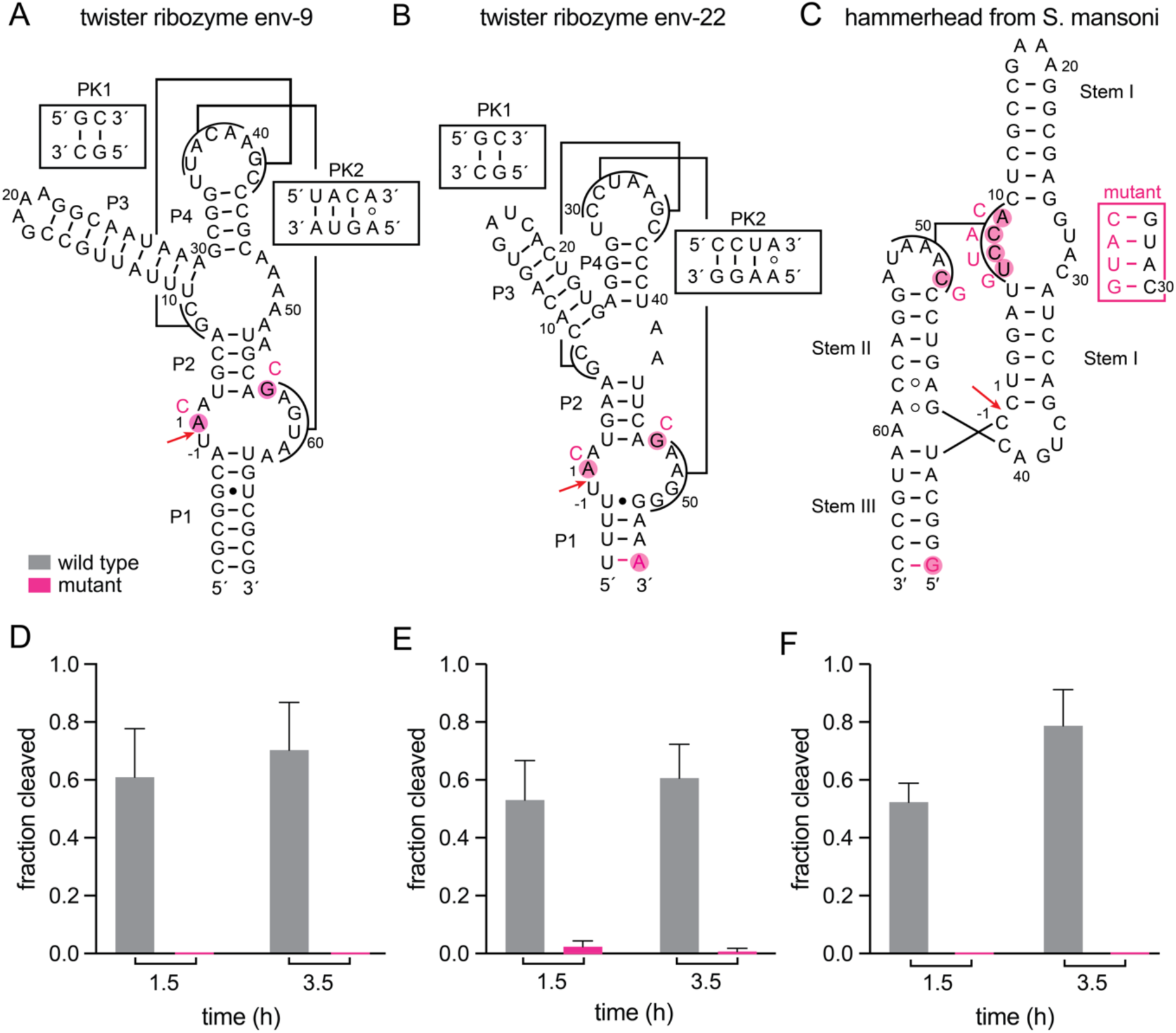
CHiTA allows for the accurate detection of *f*_cleaved_ for well-characterized active and inactive ribozymes. Secondary structure of (A) twister ribozyme env-9 (17) (B) twister ribozyme env-22 (17), and (C) a hammerhead ribozyme from *S. mansoni* (75). The cleavage site of each ribozyme is denoted with a red arrow, and tertiary interactions are presented. Bases mutated to make the inactive variant are colored magenta. In (A) and (B), the inactivating mutations target the general acid (A1) and general base (G57, G47), while in (C) the inactivating mutations prevent the formation of tertiary interactions. (D-F) Average *f*_cleaved_ for the active (gray) and inactive (magenta) variants of (D) twister ribozyme env-9, (E) twister ribozyme env-22, and (F) a hammerhead ribozyme from *S. mansoni* after 1.5 and 3.5 h of transcription. The *f*_cleaved_ shown for each ribozyme is the average of three replicates, and the error bars are the standard deviation.

All eight ribozymes were ordered as part of a single oligo pool and tested in triplicate using CHiTA. Ribozymes were transcribed for 3.5 h, with timepoints removed at 1.5 and 3.5 h. Libraries were then prepared and sequenced (see Materials and Methods). Following, reads were filtered, counted, background-corrected, and *f*_cleaved_ was calculated according to Equation 5. Replicates showed strong correlations (Figure S6).

After 1.5 h of transcription, the active env-9 twister ribozyme had an average *f*_cleaved_ of 0.61, which rose to 0.70 at 3.5 h (Figure 2D). Contrastingly, the inactive env-9 mutant did not cleave appreciably, even after 3.5 h. Similar results were obtained for the env-22 ribozyme (Figure 2E), which displayed 26- and 61-fold enhancements of the active over the inactive variant at 1.5 and 3.5 h, respectively. The hammerhead ribozyme also demonstrated a large difference in cleavage between the active and inactive variants, as the active ribozyme had an average *f*_cleaved_ of 0.79 by 3.5 h, whereas the inactive mutant had no detectable cleavage (Figure 2F). As expected, neither the wild-type CPEB3 HDV-like ribozyme with its inhibitory upstream flank nor the C57U mutant had an average *f*_cleaved_ above 0.03 (Figure S5B). These results signify that CHiTA can differentiate between active and inactive ribozymes from diverse classes.

Next, we sought to determine if cleavage was site-specific. To accomplish this, we calculated the fraction of reads at positions -3, -2, -1, 1 (the cleavage site), 2, 3, and 4 for the active and inactive ribozymes; the CPEB3 ribozyme was not included in this analysis because it was employed as a negative control. We anticipated that for the active ribozymes, the fraction of reads would be much higher at the cleavage site. If a ribozyme were miscleaving, however, the fraction of reads would be higher at a neighboring location. As expected for the active variants, the fraction of reads was much higher at the cleavage site than any of the six neighboring locations (Figure S7A, C). Moreover, for the inactive mutants, the fraction of reads at the cleavage site was smaller or similar to that of the six adjacent sites (Figure S7B, D). Thus, ribozyme self-cleavage was site-specific, for the active variants, as determined by CHiTA.

### Characterization of twister ribozymes from over a thousand diverse organisms using CHiTA

Expanding from our small-scale validation experiments, we next applied CHiTA on a large-scale to characterize 2, 625 putative twister ribozymes. Upon their initial discovery in 2014, Breaker and colleagues predicted 1, 613 unique type-P1 twister ribozymes (17). Of these, however, only 14 (<1%) were experimentally validated (17), and subsequent HT mutational studies have focused on just two representative twister ribozymes (27, 28, 31). Since 2014, additional twisters have been predicted (24, 38, 50, 65), suggesting that there may be large numbers of undiscovered variants that could aid in defining the structural and catalytic properties of twister ribozymes. Utilizing sequence similarity (Figure S1), we discovered 1, 012 new twister ribozyme candidates. As such, we sought to characterize these 2, 625 putative twister ribozymes using CHiTA.

First, the 1, 613 previously identified twister ribozyme candidates (17) and the 1, 012 novel variants that we identified were ordered as separate oligo pools, termed “literature” and “novel” oligo pools, respectively. Both oligo pools were tested for self-cleavage activity and specificity in triplicate using CHiTA. Transcription reactions were terminated after 3 h to provide sufficient time for active ribozymes to self-cleave and adequate material for sequencing.

During the computational analysis, 677 ribozymes were removed in total (Figure 3A), most because they generated the same 3′-cleavage fragment. Contrastingly, only 167 ribozymes were excluded because they had <100 reads. The remaining 1, 948 ribozymes were scored and all replicates showed robust correlations (Figure S8A-B). Values of *f*_cleaved_ ranged from 0.0-1.0 (Figure 3B), which was similar the distribution for the literature and novel oligo pools (Figure S8C-D), with average and median values of 0.76 and 0.85, respectively.

**Figure 3.**
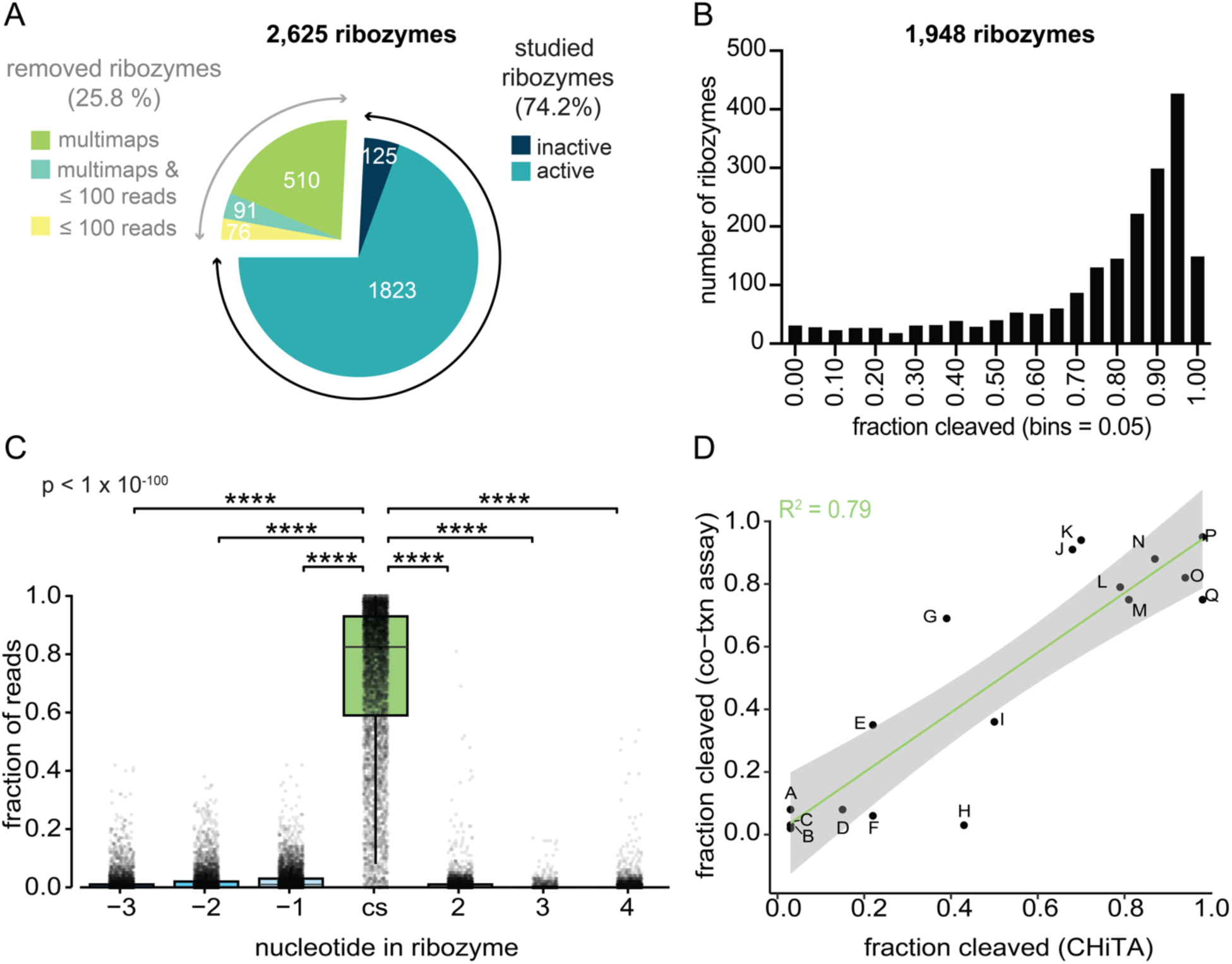
CHiTA allows for the direct detection of nearly 2, 000 twister ribozymes. (A) Pie chart depicting the number of ribozymes removed by different filters. The numbers of inactive ribozymes (*f*_cleaved_ ≤ 0.20 after 3 h of transcription) and active ribozymes (*f*_cleaved_ > 0.20 after 3 h of transcription) are provided. (B) Distribution of *f*_cleaved_ for all 1, 948 ribozymes studied (bin width = 0.05). (C) Boxplots of *f*_cleaved_ at different positions in the ribozyme relative to the cleavage site for the large-scale oligo pools. Each dot provides the average *f*_cleaved_ from triplicate experiments for a single ribozyme. Significance (p) from Kruskall-Wallis one-way ANOVA test for this analysis is listed in the upper lefthand corner. For specific comparisons, Games-Howell post-hoc tests were used on the raw data, as opposed to the average (****: p ≤ 0.0001). p-values are provided in Table S3. (D) Correlation of average *f*_cleaved_ values for 17 twister ribozymes measured using co-transcriptional cleavage assays versus using CHiTA. The line of best fit and 95% confidence interval are depicted as a green line and gray shaded area, respectively. The goodness of fit (R^2^) is provided in the upper lefthand corner. The letters correspond to the gel images in Figure S10, which were used to calculate fraction cleaved from co-transcriptional cleavage assays.

Similar to the small-scale dataset, we determined if cleavage was site-specific. Indeed, the fraction of reads was significantly higher at the cleavage site compared to any of the three sites upstream or downstream of the expected cleavage site (Figure 3C, Table S3). In general, for nucleotides upstream of the cleavage site, the fraction of upstream reads decreased as the fraction of cleavage site reads increased (Figure S9A), reflective of the specificity of cleavage. This pattern for nucleotides downstream of the cleavage site was slightly weaker (Figure S9B). Overall, these data are indicative of site-specific self-cleavage events in thousands of natural twister ribozymes as analyzed by CHiTA.

To further support calculations made with CHiTA, we selected 17 twister ribozymes from the large-scale oligo pools to test individually using a traditional gel-based approach. Because twister ribozymes cleave as they are being transcribed (17), we conducted these cleavage assays co-transcriptionally, under the same conditions as those used with CHiTA, collecting time points out to 3 h of transcription (Figure S10). For each ribozyme, we plotted the average *f*_cleaved_ at 3 h of transcription as measured by the co-transcriptional cleavage assays versus measured with CHiTA (Figure 3D). The data exemplified a goodness of fit (R^2^) of 0.79, suggesting that CHiTA can accurately assess the *f*_cleaved_ of twister ribozymes on a large-scale.

### Structural influence of imperfections in core elements of twister ribozymes

The majority of the twister ribozymes that we studied had imperfections in their secondary and tertiary structure. To discern the tolerance of imperfections within twisters, which could better inform on their structural properties and aid in discovery, we analyzed the influence of imperfections on ribozyme self-cleavage. First, sequences from the large-scale oligo pools were aligned to the consensus structural model (Figure 4A), which was composed of three interconnected stem-loops (P1/L1, P2/L2, P4/L4) and two pseudoknots (PK1, PK2) (17). Sequences were then divided into their respective structural components (P1, L1, P2, L2, PK1, P4, L4, and PK2) and imperfections within each component were identified using a custom R script. The type of imperfections identified herein included bulges (insertions into paired elements), deletions (deletions of conserved residues or base pairs), insertions (insertions into loops or pseudoknots), mismatches (non-Watson Crick Franklin base pairs in paired elements), mutations (substitutions at conserved residues), and overhangs (single-stranded nucleotides at the termini of P1). Most of the core structural elements in individual ribozymes tested here had fewer than 3 of these imperfections (Figure 4B).

**Figure 4.**
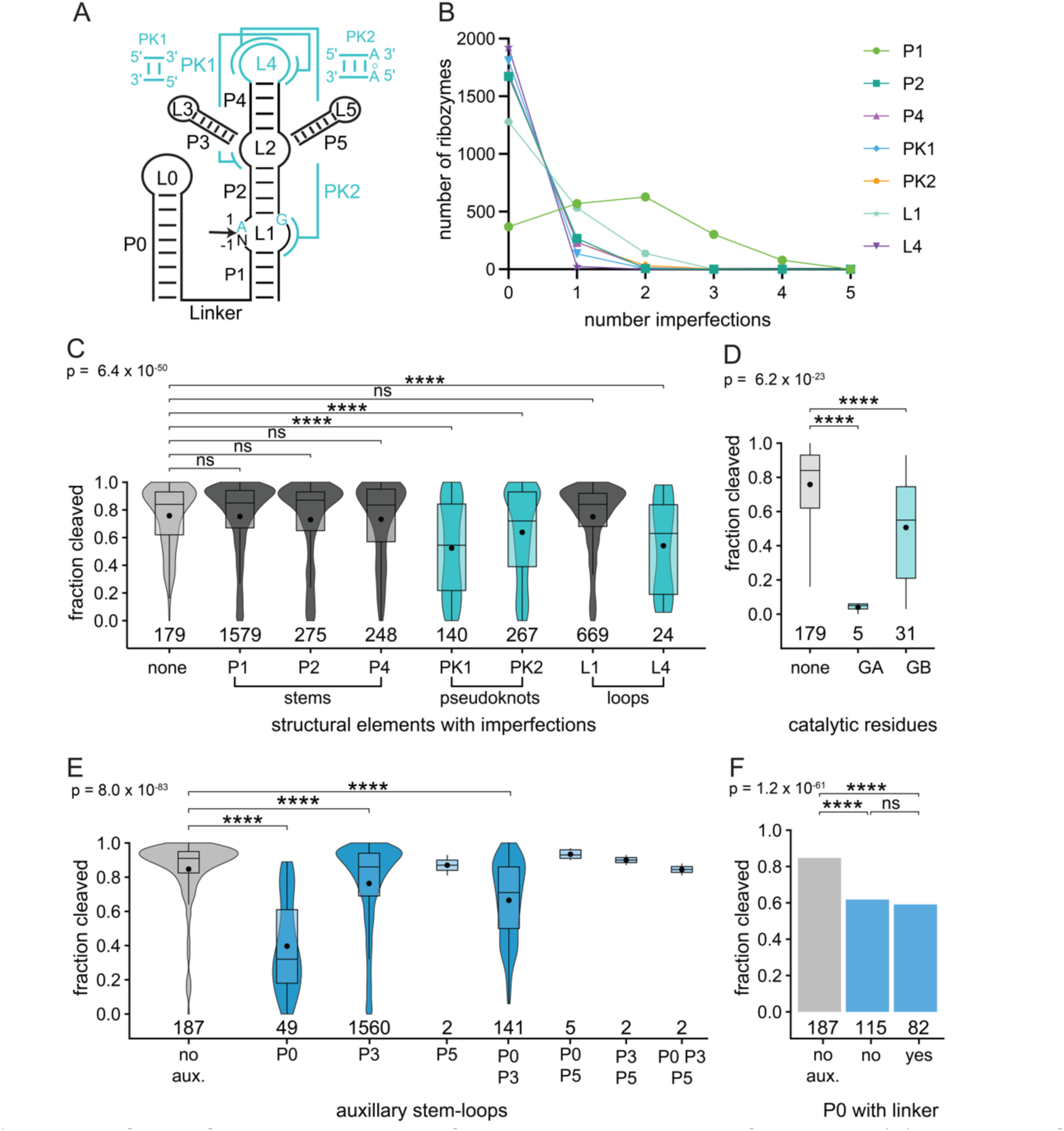
Impact of imperfections in the core of twister ribozymes on self-cleavage. (A) Depiction of the secondary structure of twister ribozymes including auxiliary stem-loops P0/L0, P3/L3, and P5/L5. (B) Line graph of the number of ribozymes with imperfections in each core element. C) Box and violin plots of the average and median *f*_cleaved_ for ribozymes with no imperfections (gray) compared to ribozymes that have at least one imperfection in one of the other listed structural elements. Note that L2 is not shown because there are no known imperfections that can occur in it. Black and teal represent structural imperfections that did not change or that reduced ribozyme self-cleavage, respectively. (D) Box plots of the average and median *f*_cleaved_ for twister ribozymes with no imperfections and those with either a mutated general acid (GA) or general base (GB). Coloring is the same as that used in panel B. (E) Effects of auxiliary stem-loops. Box and violin plots of the average and median *f*_cleaved_ for twister ribozymes with no (gray) or different combinations of auxiliary stem-loops (blue). (F) Impact of a linker sequence with P0/L0. Bar graph of the average *f*_cleaved_ for twister ribozymes with no auxiliary stem-loops (gray) or a P0/L0 with or without a linker (blue). (C-F) The black dot signifies the average *f*_cleaved_ for each group. The number of ribozymes in each category is listed below each column. Significance (p) from Kruskall-Wallis one-way ANOVA tests is provided in the upper lefthand corner of each graph. p-values are only shown for groups with ≥ 10 ribozymes to avoid small number bias. For specific comparisons, Games-Howell post-hoc tests were used (ns: not significant; *: p ≤ 0.05, **: p ≤ 0.01, ***: p ≤ 0.001, ****: p ≤ 0.0001) and values are listed in Table S4.

The number, location, and type of imperfections, as well as the length and predicted ΔG°_37_ (paired elements only) of each element, were compared against the average *f*_cleaved_. We first considered imperfections in P1. Twister ribozymes with imperfections in their P1 stem (Figure S11A) did not exhibit a significant reduction in their average *f*_cleaved_ relative to ribozymes with no imperfections (control group) (Figure 4C). This suggested imperfections in P1 were well tolerated. Moreover, no relationship was found between the average *f*_cleaved_ and the length (Figure S11B-C) or the predicted ΔG°_37_ (Figure S11D) of P1. Additionally, the average *f*_cleaved_ appeared to be largely independent of the number (Figure S11E), location (Figure S11F), and type of imperfections (Figure S11G-H). Indeed, ribozymes with a bulged P1 displayed an *elevated* average *f*_cleaved_ of 0.86 relative to the average *f*_cleaved_ of 0.76 for ribozymes with no imperfections, albeit the sample size of bulged helices was relatively small at only n=16 (Figure S11G). Consequently, twister ribozymes were essentially impervious to imperfections in their P1 stem, which is consistent with previous studies that demonstrated that P1 is dispensable (80).

Similar to P1, imperfections in the P2 stem of twister ribozymes (Figure S12A) were well tolerated (Figure 4C). Once again, no relationship was identified between the average *f*_cleaved_ and the length (Figure S12B-C) or predicted ΔG°_37_ (Figure S12D) of P2. As with P1, the average *f*_cleaved_ had little dependence on the number (Figure S12E), location (Figure S12F), general (Figure S12G), or specific type (Figure S12H) of imperfections in P2, nor on the combination of type and location of imperfections (Figure S12I). Thus, twister ribozymes were largely unaffected by imperfections in their P2 stem.

Akin to P1 and P2, imperfections in P4 (Figure S13A) were highly tolerated by twister ribozymes (Figure 4C). The length (Figure S13B-C) as well as the predicted ΔG°_37_ (Figure S13D) of P4 showed no correlation with the average *f*_cleaved_. For ribozymes with imperfections in their P4, the average *f*_cleaved_ was not influenced by the number of imperfections (Figure S13E). However, imperfections at the fourth base pair of P4 did not tolerate imperfections as well (Figure S13F); twister ribozymes with an imperfection at this location yielded a reduced average *f*_cleaved_ of 0.58 relative to the average *f*_cleaved_ of 0.76 for ribozymes with no imperfections. Due to its proximity to the apical loop, imperfections at location 4 of P4 may disrupt orientation of PK1 and PK2, thereby inhibiting self-cleavage. Both deletiosn and mismatches in P4 were well tolerated (Figure S13G). Further examination of the specific type of imperfections demonstrated no effect on the average *f*_cleaved_, although A·C mismatches lowered the average *f*_cleaved_ by 0.14 compared to the control group (Figure S13H). When analyzed in conjunction with location, A·C mismatches at location 4 had a slightly reduced average *f*_cleaved_ of 0.60 (Figure S13I). Therefore, twister ribozymes did tolerate imperfections in P4, except for at location 4.

While twister ribozymes largely tolerated imperfections in their core stems, some imperfections in PK1 (Figure S14A) severely diminished the average *f*_cleaved_ (Figure 4C). The number of imperfections in PK1 influenced *f*_cleaved_, as one imperfection reduced the average *f*_cleaved_ from 0.76 to 0.53, while two decreased it further to 0.36 (Figure S14B). Imperfections in each of the two locations produced a significantly lower average *f*_cleaved_ (Figure S14C). For the specific type of imperfections, *f*_cleaved_ remained relatively unaltered by G·U and U·G wobbles (Figure S14D); one ribozyme with a G·U and U·G wobble in its PK1 had an average *f*_cleaved_ of 0.82 (Figure S14B). However, the average *f*_cleaved_ was greatly diminished for most other mismatches (Figure S14D), leading to a bimodal distribution as seen in Figure 4C. These findings are further substantiated by evaluation of the type and location in combination (Figure S14E). In sum, twister ribozymes were sensitive to imperfections in PK1 and the extent of sensitivity was dependent upon the number and type of imperfection.

Imperfections in PK2 (Figure S15A) also reduced the average *f*_cleaved_ relative to ribozymes with no imperfections, but not as much as imperfections in PK1 did (Figure 4C, Table S4). As seen with other core elements, the average *f*_cleaved_ was not contingent upon either the length (Figure S15B-C) or predicted ΔG°_37_ (Figure S15D) of PK2. The number of imperfections, however, did correlate with *f*_cleaved_, as one, two, or three imperfections reduced the average *f*_cleaved_ from 0.76 for the control group to 0.65, 0.55, and 0.28, respectively (Figure S15E). Apart from ribozymes with two or more imperfections in their PK2, locations 2 and 3 were more susceptible to imperfections, as their average *f*_cleaved_ dropped from 0.76 for the control group to 0.63 and 0.57, respectively (Figure S15F). It may be that imperfections in the middle of pseudoknots were more effective at disrupting base pairing (81). Broadly, many mismatches in PK2 were not well-tolerated, as the average *f*_cleaved_ varied significantly relative to ribozymes with no imperfections (Figure S15G). However, A·G, C·A, C·C, and U·G mismatches were better tolerated (Figure 15H-I). Interestingly, A-to-C mutations at the known *trans*-Watson Crick Franklin A·A interaction (63) displayed no statistically significant difference compared to the control group, while a mutation to a G did (Figure S15H). Therefore, the impact of imperfections within PK2 was dependent upon the number, location, and type of imperfection, similar to PK1. However, PK2 better tolerated weaker base pairing than PK1 because it has two additional base pairs.

Broadly, L1 (Figure S16A) exemplified some tolerance of imperfections (Figure 4C). The length of the 5′-and 3′-strands of L1 varied greatly and did not correlate with the average *f*_cleaved_, therefore being relatively tolerant of insertions (Figure 16B-C). A single imperfection in L1 did not significantly impact the average *f*_cleaved_, while two imperfections reduced it only slightly, from 0.76 to 0.67 (Figure 16D). Imperfections upstream of the cleavage site (5′_-1) or downstream of PK2 (3′_6) were broadly tolerated (Figure 16E). In contrast, imperfections downstream of the cleavage site (5′_2, 5′_3), which were relatively few, demonstrated an average *f*_cleaved_ of just 0.19 across all locations (Figure S16E). Regarding the general type of imperfection, insertions had little effect on the average *f*_cleaved_, but mutations were pernicious lowering it to 0.15 (Figure S16F). Specifically, we considered the effect of mutations at four key locations. Nucleotide -1 was canonically represented as a U (17), but in our oligo pools, all four nucleobases were relatively common (ranging from 4 to 8.5%). All nucleobases at n-1 were tolerated, albeit A and G exhibited a slightly lower average *f*_cleaved_ of 0.72 and 0.68, respectively (Figure S16G). Conversely, mutation of either catalytic residue was deleterious (Figure 4D). Substitution of A with U or C at the general acid (5’_1) completely abolished activity for all 5 ribozymes (Figure 16H). Substitution of G with A at the general base (3’_1) partially reduced the average *f*_cleaved_ from 0.76 to 0.54 and changes to a C or U furthered lowered the average *f*_cleaved_ to 0.37 and 0.34, respectively (Figure 16I). Similarly, mutations at A2 resulted in near loss in activity (Figure S16J). Note, that while mutations to conserved residues were rather scarce in these oligo pools (Figure 4D, S16I), additional examples may occur in nature. In sum, L1 tolerated imperfections at its base, but was not accepting of changes downstream of the cleavage site.

Unlike other core structural elements, L2 (Figure S17A) was a highly variable region according to the original consensus model (17). Instead of analyzing imperfections, we determined how the length of L2 impacted the extent of ribozyme self-cleavage, excluding P3/L3 and P5/L5 from this analysis. Both the 5′- and 3′-strands of L2 displayed a wide range of average *f*_cleaved_ values that depicted no direct correlation with length (Figure S17B-C), suggesting L2 was highly amenable to different insertions on either strand.

Finally, some imperfections in L4 (Figure S18A) significantly lowered the average *f*_cleaved_ (Figure 4C). We limited analysis of L4 to only those nucleotides that did not participate in pseudoknots because these were analyzed above in PK1 and PK2. As with the other core structural elements, the average *f*_cleaved_ had no dependence upon the length of L4 (Figure S18B). Imperfections at locations 1 and 9 were well tolerated, but changes at location 6, which were also the most prevalent, demonstrated a significantly reduced average *f*_cleaved_ of 0.42 relative to the control group value of 0.76 (Figure S18C). Both insertions and mutations reduced the average *f*_cleaved_ relative to ribozymes with no imperfections (Figure S18D). More specifically, though, A6G mutations drastically reduced the average *f*_cleaved_ to 0.15, while A6U mutations did not (Figure S18E). This resulted in the bimodal distribution observed in Figure 4C and S18D. Alignment of these six A6G L4 mutant twisters showed they were highly similar (Figure S19) and were not predicted to assume the consensus structure of twister ribozymes when folded with *Fold* (55), suggesting that they may misfold.

In sum, twister ribozymes tolerated imperfections throughout their core elements. They were fully tolerant of imperfections in P1, P2, and P4, wobbles in PK1 and especially PK2, and imperfections in L1 and L4, albeit in a positional fashion.

### Structural influence of auxiliary stem-loops on twister ribozyme self-cleavage

Twister ribozymes sometimes possess one or more auxiliary stem-loops, whose impact on self-cleavage has not yet been elucidated (17, 63, 78). To determine how auxiliary stem-loops influence the extent of self-cleavage, we compared the average *f*_cleaved_ for ribozymes with and without a P0/L0, P3/L3, or P5/L5 stem-loop, including any combination thereof. No imperfections were noted for any auxiliary stem-loops as the extent of base pairing could not always be accurately determined. However, the length and predicted ΔG°_37_ were calculated for each auxiliary stem-loop and their correlations with *f*_cleaved_ were examined.

The P0/L0 stem-loop was located immediately upstream of the 5′-terminus of twister ribozymes (Figure S20A) (17). Of the candidates we analyzed, P0/L0 occurred in approximately 10% of the ribozymes. In our oligo pools. Compared to ribozymes with no auxiliary stem-loops, which had an average *f*_cleaved_ of 0.85, ribozymes with a P0/L0 exhibited a much lower extent of cleavage, with an average *f*_cleaved_ of just 0.40 (Figure 4E). This reduced self-cleavage may be reflective of inhibitory interactions between P0/L0 and the ribozyme that elicit an alternate, inactive conformation of the ribozyme. No relationship was found between the average *f*_cleaved_ and the length or predicted ΔG°_37_ of P0/L0 (Figure S20A). For 82 twisters, a linker sequence separated P0/L0 and the ribozyme (Figure S20B). We postulated that this linker might limit interactions between P0/L0 and the ribozyme. However, its presence did not influence the average *f*_cleaved_ relative to ribozymes with a P0/L0 but no linker (Figure 4F). Similarly, the length of the linker did not substantially influence the average *f*_cleaved_, although some ribozymes with a longer linker displayed a higher extent of self-cleavage (Figure S20B).

Of the candidates we analyzed, 1705 twisters had a P3/L3 stem-loop, found on the 5’-side of L2 (Figure S20C). On average, twister ribozymes with a P3/L3 exhibited a somewhat lower average *f*_cleaved_ of 0.76 relative to twisters with no auxiliary stem-loops, which had a *f*_cleaved_ of 0.85 (Figure 4E). Combinations of P3 and P0 presented an intermediate average *f*_cleaved_ of 0.67, reflective of the inhibitory influence of P0/L0. Analogous to P0/L0, there was no correlation between the average *f*_cleaved_ and the length or predicted ΔG°_37_ for P3/L3, although more ribozymes with a shorter P3/L3 or higher ΔG°_37_ achieved a higher average *f*_cleaved_ (Figure S20C). Notably, some ribozymes with a P3/L3 longer than 50 nt had an average *f*_cleaved_ greater than 0.40 (Figure S20C). Therefore, P3/L3 generally did not interrupt self-cleavage, and active ribozymes (*f*_cleaved_ > 0.20) could tolerate long P3 hairpins.

Twister ribozymes with a P5/L5 stem-loop, found on the 3′-side of L2 (Figure S20D), were not as common in our library and mostly found in combination with P0/L0, P3/L3, or both. The range of lengths and predicted ΔG°_37_ for P5/L5 was not dissimilar to that for P3/L3. Curiously, every twister with a P5/L5 had an average *f*_cleaved_ greater 0.80 (Figure 4E). Again, no relationship between *f*_cleaved_ and the length or predicted ΔG°_37_ for P5/L5 (Figure S20D) existed. It appeared that twister ribozymes were very tolerant of P5 stem-loops, although the sample size analyzed here was small.

### Structural influence of total imperfections on twister ribozyme self-cleavage

Broadening our analysis, we next assessed how imperfections within the full-length ribozyme impacted self-cleavage. First, we employed our custom R script to determine the length and predicted ΔG°_37_ for full-length ribozymes. As previously evidenced with the individual components, no relationship was evident between the average *f*_cleaved_ and the length (Figure S21A) or the predicted ΔG°_37_ for each ribozyme (Figure S21B). However, the length and predicted ΔG°_37_ (Figure S21C) were weakly correlated with each other, suggesting longer regions are pairing.

Next, we investigated how imperfections within one, two, or three different structural elements contributed to self-cleavage (Figure 5). Utilizing our custom R script, we identified combinations of structural elements with imperfections. Due to previous studies that have established that P1 is dispensable (78), which is also supported by our structural analysis, it was not considered an imperfection here. Ribozymes with one or two imperfect elements were plotted as a heat map, where all ribozymes with a single imperfect element aligned on the diagonal and those with two imperfect elements fell below the diagonal (Figure 5A). Imperfections at L1, P2, P4, or L4 were generally well accommodated, having an average *f*_cleaved_ > 0.50, and combinations of imperfections of these elements with imperfect PK1, PK2, or the general base were also tolerated (Figure 5A). In contrast, ribozymes with imperfections at the general acid, either alone or in combination with other imperfect elements, were almost always detrimental to self-cleavage (Figure 5A). Ribozymes with three imperfect structural elements were relatively rare in our oligo pools and had a range of average *f*_cleaved_ from 0.06 to 0.58 (Figure 5B). Most of these ribozymes had imperfections in PK1, PK2, or at their general base, which were tolerated when matched with imperfections in L1, P2, or P4, but not when matched with each other (Figure 5B). In summary, twister ribozymes supported multiple imperfect elements, except when an imperfection occurred at both the pseudoknots and catalytic residues.

**Figure 5.**
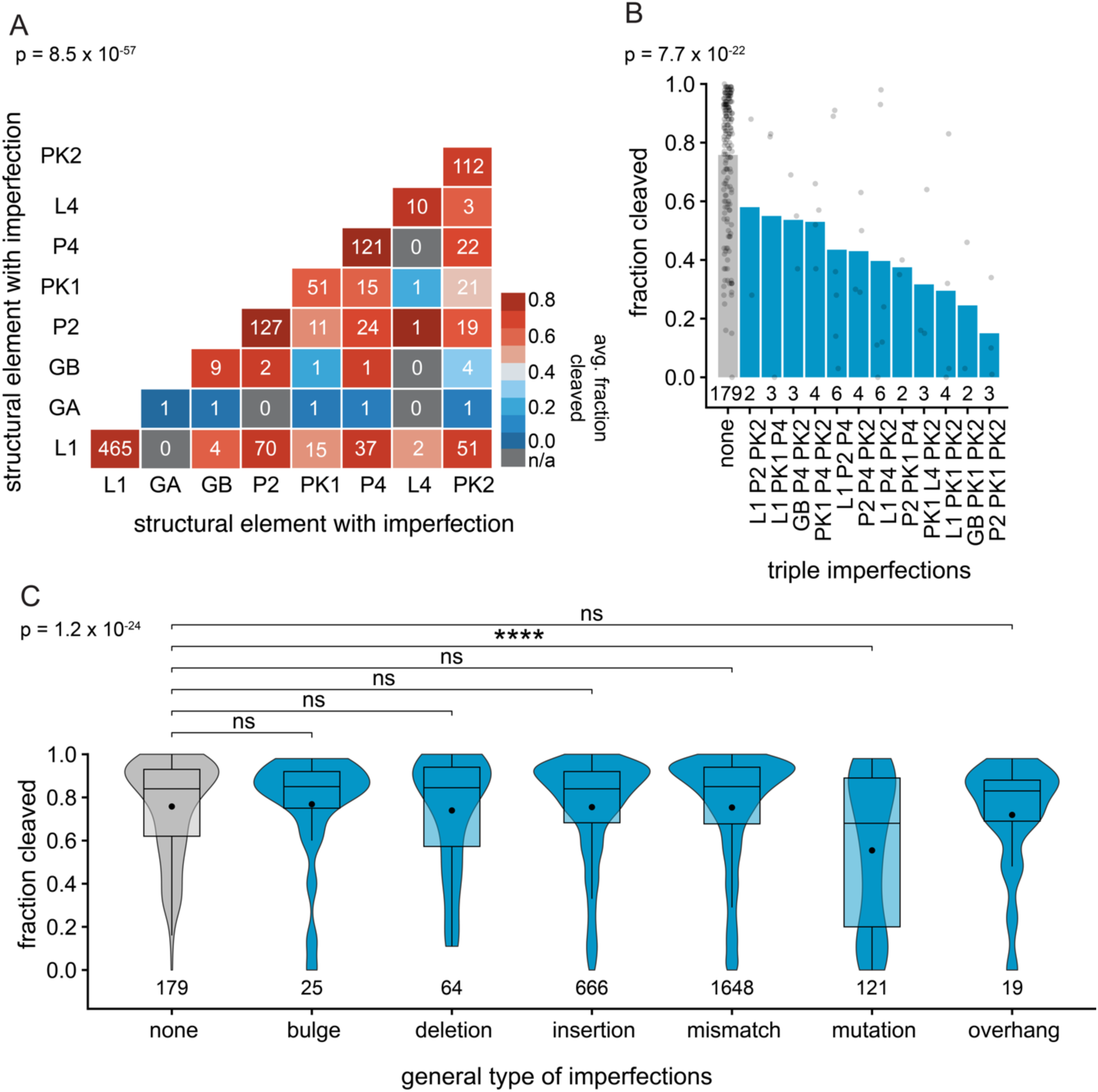
Impact of the number and type of imperfections in twister ribozymes on self-cleavage. (A) Heat map of ribozymes with one or two structural elements that contain imperfections. GA and GB stand for general acid and general base, respectively. Because the P1 stem has no impact on ribozyme self-cleavage it was ignored. The color corresponds to the average *f*_cleaved_ for each group. (B) Bar graph of the average *f*_cleaved_ for ribozymes with three structural elements that contain imperfections. (C) Box and violin plot of the average (black dot) and median *f*_cleaved_ for the general types of imperfections found in all ribozymes (blue) compared against ribozymes with no imperfections (gray). (B-C) The number of ribozymes in each category is listed below each graph. Significance (p) from Kruskall-Wallis one-way ANOVA tests is provided in the upper lefthand corner of each graph. For specific comparisons Games-Howell post-hoc tests were used (ns: not significant; *: p ≤ 0.05, **: p ≤ 0.01, ***: p ≤ 0.001, ****: p ≤ 0.0001) and values are provided in Table S12.

We then determined how the general type and number of imperfections influenced ribozyme behavior. Bulges, deletions, insertions, mismatches, and overhangs were generally well tolerated, as all had a similar average *f*_cleaved_ to ribozymes with no imperfections (Figure 5C). This was further substantiated by lack of dependence of the average *f*_cleaved_ on the number of deleted or inserted nucleotides (Figure S22A, S22B) or the length of the overhang (Figure S22C). Mismatches were the most common type of imperfection, and up to 4 were generally tolerated, while 5 and 6 mismatches significantly reduced the average *f*_cleaved_ from 0.76 to 0.63 and 0.47, respectively (Figure S22D). Contrastingly, mutations reduced the average *f*_cleaved_ from 0.76 to 0.57 and having two mutations reduced it further to 0.10 (Figure S22E). More generally, though, we observed no correlation between the total number of imperfections within twister ribozymes and the average *f*_cleaved_ (Figure S22F). Overall, self-cleavage was most dependent upon the type and number of mutations, as opposed to other types of imperfections.

### Twister ribozymes are found in many organisms

Upon their discovery in 2014, the Breaker laboratory predicted twister ribozymes in 23 different organisms including anaerobic bacteria, cnidarians, insects, and plants as well as a fungus, flatworm, fish, and a virus (Figure S23A, inset) (17). Additional candidates were unearthed in plants, fish, flatworms, gastropods, and reptiles by Rfam (50) and Liu et al. (65), while Lee et al. detected over 3, 500 twister ribozymes from metatranscriptomes (24). In total, twister ribozymes have been predicted in 78 different organisms, as well as in environmental samples and metatranscriptomes (17, 24, 50, 65).

To distinguish our search, we sought to expand the number of organisms with twister ribozymes and so surveyed a wider array of organisms (Figure S23A, column 2). Emphasis was placed on groups with few to no current candidates, including amphibians, birds, and reptiles. Through our survey, we added 1, 129 new species that encompassed new groups such as amphibians, arachnids, arthropods, bivalves, bryozoans, crustaceans, echinoderms, isopods, rotifers, and various types of worms. Collectively, we enriched the pool of organisms predicted to possess twister ribozymes by over 14-fold (Figure S23A, column 3).

Of the candidates tested from the large-scale experiments, most of the active ribozymes originated from environmental samples, fish, flatworms, insects, and plants (Figure 6A, Table S14). Interestingly, all of the plant species with active twister ribozymes belonged to flowering plants and no candidates were initially detected in mosses, algae, or the *Brassicaceae* family. While those active ribozymes in plants came from hundreds of different species, the ones in flatworms resided primarily in *S. mansoni* and related species (Figure S23B). Notably, this was the first time intrinsically active twister ribozymes were identified within reptiles and amphibians (Figure S23B). After our experimental analysis no active twister ribozymes were found in birds and cephalopods (Figure 6A, S23C). Multiple inactive candidates were also in fungi and reptiles (Figure 6A, S23C). Thus, while our survey drastically expanded the number of species with intrinsically active twister ribozymes, many types of organisms, most notably mammals, still lacked any strong candidates.

**Figure 6.**
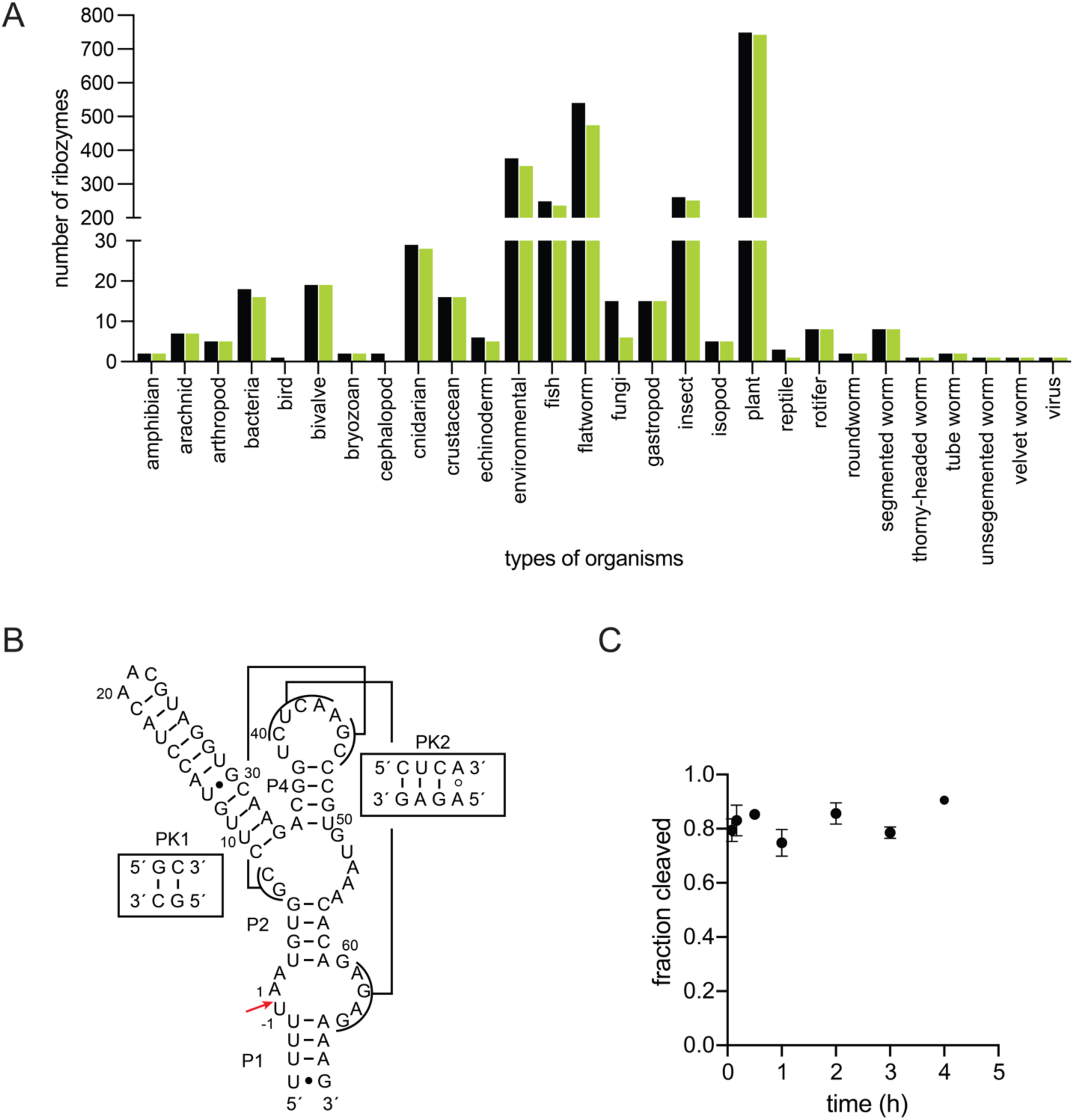
Twister ribozymes are found in many organisms, including mammals. (A) Bar graph of the types of organisms with predicted (black) and active (green) ribozymes studied using CHiTA. Active ribozymes are those defined as having an average *f*_cleaved_ > 0.20. (B) Secondary structure of Ttru-1-1, an intrinsically active twister ribozyme found in *Tursiops truncatus* (bottlenose dolphin). The cleavage site of the ribozyme is denoted with a red arrow, and tertiary interactions are presented. (C) Scatter plot of the *f*_cleaved_ vs time for the Ttru-1-1 from co-transcriptional cleavage assays tested in triplicate.

To address whether ribozymes might exist in such organisms, we conducted another computational search using updated structural descriptors. Utilizing the active twister ribozymes studied with CHiTA, we constructed updated consensus sequences (Figure S24) and structural descriptors (Figure S4) for twister ribozymes without a P3/L3 or P5/L5 (Figure S24, S4A), a P3/L3 (Figure S24B, S4B), and a P5/L5 (Figure S24C, S4C); these descriptors permitted imperfections, as well as loosened the base pairing rules for stems, as appropriate. Then, the descriptors and select genomes were input into our Computational Discovery Pipeline, which had three parts: 1. search the input genome(s) for sequences matching the provided descriptor with *RNArobo* (23), 2. align sequences to the consensus structure with *cmalign* from the *Infernal* package (22), and 3. fold the candidates using *R2R* (54). To minimize the number of false positives, sequences with a bit-score below zero were culled.

We benchmarked this pipeline for its ability to identify the known active twister ribozymes from the large-scale oligo pools. For ribozymes without a P3/L3 or P5/L5, 132 out of 217 were correctly identified (Figure S24D, Table S15). Similarly, for ribozymes with a P3/L3 72% of all active candidates were detected, but only 26% of candidates with a P5/L5 were found (Figure S24D, Table S15). In all, 71% of all active candidates were correctly identified (Figure S24D, Table S15).

The pipeline was then applied to search for additional twister ribozymes, focusing on 54 organisms with no known twisters. Our search centered primarily around mammals, green algae, and moss since these groups were not represented in the large-scale oligo pool. Across all these organisms, 8, 665 candidate ribozymes were identified (Table S16). From these, we selected three candidates to test using co-transcriptional cleavage assays: two from *Homo sapiens* (human) Hsap-1-1 (Figure S25A) and Hsap-1-2 (Figure S25B) and one from *Tursiops truncatus* (bottlenose dolphin) Ttru-1-1 (Figure 6B). Neither, Hsap-1-1 or Hsap-1-2 self-cleaved to an appreciable amount in 1 day (Figure S26A-B). However, Ttru-1-1 was highly active, with an average *f*_cleaved_ of 0.79 within 5 min (Figure 6C, S26C). Intriguingly, the sequence of Ttru-1-1 was most similar to Nvi-1-37, differing by only four nucleotides in P3 (Figure S27). In conclusion, we greatly expanded the number of organisms that possess twister ribozymes and in so doing discovered the first intrinsically active mammalian twister ribozyme.

## Discussion

Advances in bioinformatics have facilitated the prediction of tens of thousands of putative naturally occurring ribozymes. Traditional biochemical techniques analyze candidates individually, rendering such methods impractical for large-scale assessments. To circumvent this issue, we developed CHiTA, which employs MPOS and NGS in a scarless fashion, to experimentally characterize large and diverse pools of putative ribozymes. We first applied CHiTA on a small-scale to examine its capacity to differentiate active and inactive ribozymes. Expanding upon this, CHiTA was employed to quantify self-cleavage for 2, 625 naturally occurring twister ribozyme candidates, 1, 613 of which were predicted in literature (17) and 1, 012 of which we identified. A comprehensive structural analysis was then conducted to determine the impact of imperfections on ribozyme self-cleavage. This revealed a broad tolerance for imperfections in most structural elements for twisters, except catalytic residues and pseudoknots. Utilizing the information garnered from the structural analysis, we performed further bioinformatic searches and discovered the first intrinsically active twister ribozyme in mammals.

## CHiTA as a means to quantify ribozyme self-cleavage *en masse*

CHiTA provides a HT ribozyme characterization pipeline that affords several advantages over existing methods. Many HT methodologies have been developed to study all single and double mutants of a single ribozyme (26). Employment of MPOS to generate the temple DNA, however, permits exploration of the full structural diversity predicted in naturally occurring small nucleolytic ribozymes from diverse organisms. Different circular permutations, as well as ribozymes with of lengths and with different auxiliary-stem loops can be analyzed simultaneously. Further, CHiTA can be applied to test mixed pools containing different classes of ribozymes, as demonstrated here on a small-scale. Importantly, CHiTA also quantifies the intrinsic extent of self-cleavage for ribozymes without the influence of artificial flanking sequence. Inclusion of the BsaI restriction site within the reverse PCR primer binding site, allows for the scarless removal of the reverse PCR primer binding site and the BsaI site itself (Figure 1B), which offers advantages other over reported HT methods that use MPOS to study ribozymes (46, 25). Third, the scoring pipeline can be customized to suit the needs of the user. Parameters such as the read length, number of mutations, and mapping score can be adjusted in our custom python script.

As with any method, though, CHiTA faces some limitations. Regarding the Experimental Pipeline, fragmentation of the RNA backbone, aborted transcripts, and DNA side-products from the library preparation generate reads starting at different positions, which will be reflected as noise in the sequencing data. This is addressed both experimentally and computationally in CHiTA. Gel purification steps remove undigested DNA, transcription aborts, and unligated product during preparation of the sequencing libraries (Figure 1B). We also performed an on-column DNase treatment to remove template DNA prior to constructing the libraries (Figure 1B). Computationally, we background correct each ribozyme to minimize random starts (Figure 1C). Another limitation is that CHiTA cannot discern between cleaved products from different ribozymes that generate the same 3′-cleavage fragment. Finally, to minimize biases inherent to NGS, we did not analyze ribozymes with a low sequencing depth, defined here as < 100 cleaved and uncleaved reads in any replicate.

Moreover, CHiTA is versatile and can be applied to other classes of ribozymes. For instance, it could be applied to study circular permutations of hammerhead or twister ribozymes (9) as well as non-circular permutations of hairpin ribozymes (36). Further, the cleavage conditions could be altered to test how different environmental conditions influence catalysis. As such, CHiTA could prove to be an invaluable tool for screening thousands of candidate ribozymes for synthetic applications.

## Twister ribozymes tolerate many imperfections in their secondary structures

CHiTA characterized 1, 948 twister ribozymes that exhibited a range of average *f*_cleaved_ from 0.0 to 1.0 (Figure 3B). For those ribozymes that did not cleave, the lower the average *f*_cleaved_ could be due to an imperfection in the secondary structure of the ribozyme, such as a mutation to the general acid or general base (Figure 4D). Biologically, these ribozymes may not need to cleave quickly, if at all, *in vivo* to achieve their function. Some twisters that were predicted to not have any imperfections, however, also demonstrated a *f*_cleaved_ ≤ 0.20. As such, misfolding of some twister ribozymes may preclude self-cleavage.

From the active ribozymes, we generated a new consensus sequence (Figure 7A) to reflect changes revealed through our analysis. While the consensus sequence remained relatively unchanged (17, 50), base identity within helical elements became more generalized. Our comprehensive structural analysis supported this, as it was observed that paired elements P1, P2, and P4 tolerated most imperfections (Figure 4C). Previous studies have demonstrated that P1 is dispensable for ribozyme activity (80), and HT mutagenesis experiments of Osa 1-4 showed that P2 and P4 were broadly tolerant of mismatches (28, 82). Our experiments revealed that, in addition to mismatches, these core stems could accommodate bulges, deletions, and overhangs (Figure 7B). The tolerance of the core stems of twister ribozymes can be attributed to their 3D organization (63, 76) in which P1, PK2, P2, and PK1 are coaxially stacked (Figure7C). As such, they can likely project imperfections away from any sites of tertiary interactions and overcome destabilization induced by one or two imperfections with strong base pairing or stacking interactions between the helices.

**Figure 7.**
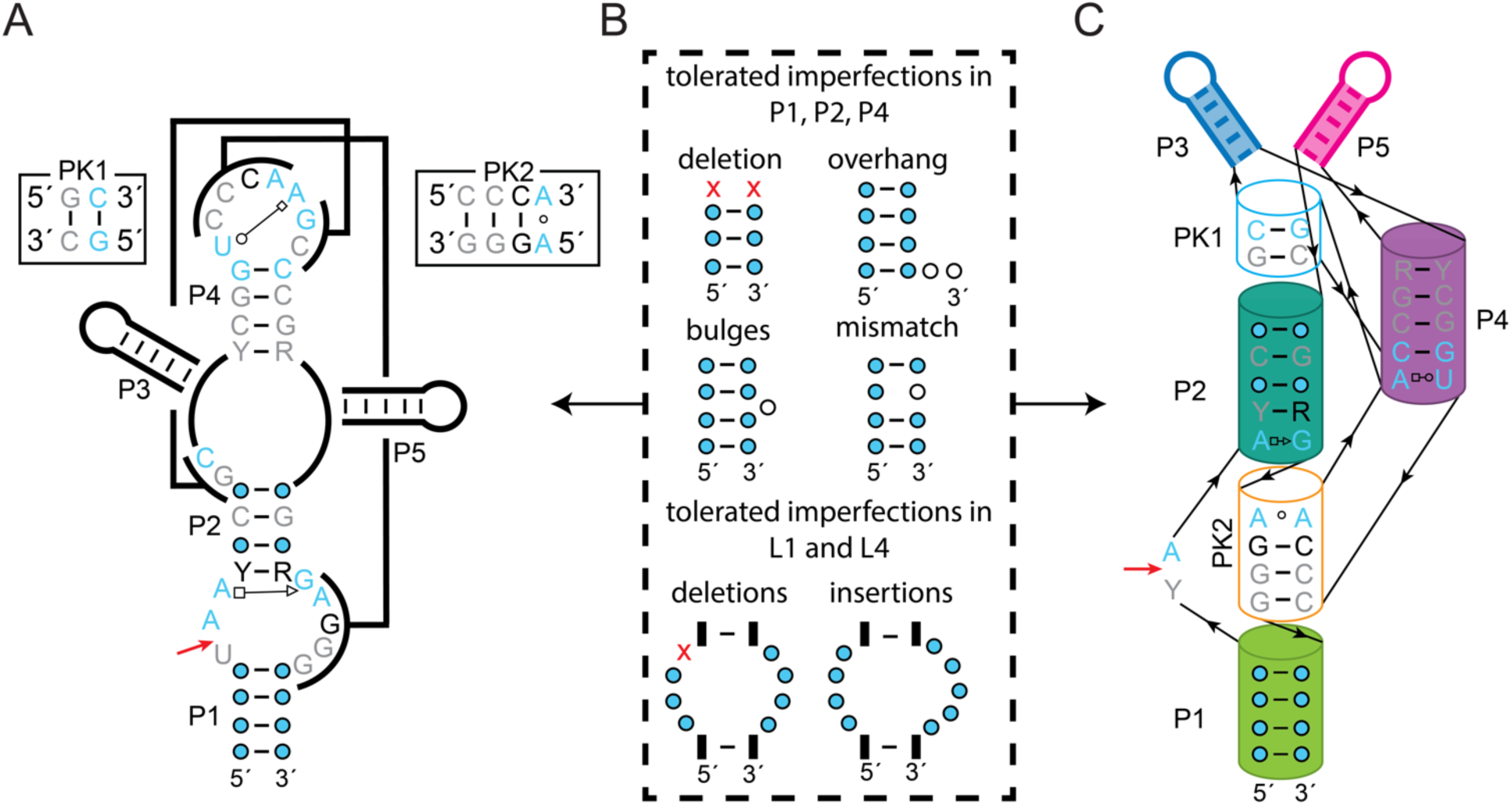
Consensus sequence of active twister ribozymes studied using CHiTA. All diagrams were made using active ribozymes (*f*_cleaved_ > 0.20). Blue, black, and gray nucleotides are conserved 97%, 90%, and 75%, respectively. The cleavage site is marked with a red arrow. (A) Consensus secondary and tertiary structure of all active twister ribozymes. (B) Examples of the type of imperfections tolerated by twister ribozymes in their paired elements and loops. (C) Consensus structure plotted onto the 3D-shape of twister ribozymes, modeled after Liu et al. (63). Paired elements are colored according to Figures S11-S18. Helical elements tolerant of imperfections are fully colored, while those susceptible to imperfections are outlined.

The loops of twister ribozymes were also somewhat tolerant of imperfections (Figure 7B). Insertions into L1 upstream of the cleavage site or downstream of PK2 had no significant impact of the average *f*_cleaved_ (Figure S16J). This is unsurprising since P1 is not important for self-cleavage (80) and the placement of these insertions in L1 may be thought of as an extension of P1. Contrastingly, imperfections at other sites within L1 most likely disrupted the orientation of the active site or tertiary interactions, thereby hindering self-cleavage. Mutations to conserved residues in L1, including the general acid and general base, were also deleterious to ribozyme activity (Figure 4D, S16F), consistent with mutagenesis studies (28, 82, 78, 63). Similarly, imperfections in L4 decreased the average *f*_cleaved_, although the extent of attenuation was dependent upon the type and location of the imperfection (Figure 4C, S18E). While these observations are supported by previous studies (28, 63, 78, 82), we found that some mutations at the sixth nucleotide of L4 exhibited different behavior than previously reported (Figure S18E). We attribute this difference to secondary characteristics of the ribozyme, rather than to the mutations themselves due to the small sample size. However, these mutations may warrant additional investigations in the future.

Tertiary interactions in PK1 and PK2 were more sensitive to imperfections (Figure 4C). This is likely because they are relatively small at two or three base pairs, and thus intolerant of weakening. For both pseudoknots the average *f*_cleaved_ was dependent upon the number and type of imperfection. In fact, the stability of mismatches in PK1 and PK2 followed the expected thermodynamic stability of non-Watson Crick Franklin base pairs, accommodating G·U wobbles most readily (83). Our findings are consistent with HT mutagenesis experiments of twister ribozymes, which demonstrated that both PK1 and PK2 were susceptible to mismatches (27, 28, 82). However, we were also able to highlight that insertions within PK2 need not be deleterious to ribozyme activity (Figure S15G). Indeed, imperfections in PK2 were somewhat less debilitating to the ribozyme than those in PK1 (Figure 4C), most likely due to its higher number of base pairing interactions as well as its placement in the crystal structure as it stacks between P1 and P2 (Figure 7C) (17, 28, 31, 63, 76). PK1 is located above P2 and occurs as the last stacked element if a P3 is not present (Figure 7C) (63, 76). It may be that an extended P3 could stabilize a two-base pair PK1 via stacking interactions.

Looking at combinations of the above structural elements revealed that imperfections in multiple components had no additional adverse effects, although in some cases the number of ribozymes was very small (Figure 5A, 5B). In fact, some ribozymes achieved a high average *f*_cleaved_ with three imperfect structural elements (Figure 5B). One unique feature of our study was the inclusion of different types of imperfections such as bulges, deletions, insertions, and overhangs; this is the first time when such imperfections were analyzed in a HT study of twister ribozymes. Broadly speaking, bulges, deletions, insertions, mismatches, and overhangs were well tolerated amongst the twister ribozymes we studied (Figure 5C).

Twister ribozymes gain further sequence and structural diversity in their P0/L0, P3/L3, and P5/L5 auxiliary stem-loops. The upstream hairpin P0/L0 reduced the average *f*_cleaved_, irrespective of its length or predicted ΔG°_37_, or presence of a linker (Figure 4E-F, S20A-B). This upstream element may interact with the ribozyme to generate inactivated conformations. Interestingly, P3/L3 and P5/L5 did not have much of an impact on the average *f*_cleaved_ (Figure 4E, S20C-D). No consensus sequence could be generated for these highly variable stem-loops and their role in twister ribozymes remains unclear.

### Twister ribozymes are harbored by many organisms

Using our structural analysis, we developed descriptors to search for additional candidate twister ribozymes with an emphasis on mammals. Prior to our study, twister ribozymes had been predicted in only 78 organisms (Figure S23A). We were able to expand this number 14-fold. Despite this, we did not find active ribozymes in groups like birds and mammals. As such, we used our findings to generate structural descriptors and developed the Computational Discovery Pipeline to identify candidate ribozymes within this group of organisms. Our search resulted in the identification of the first intrinsically active twister ribozyme from mammals in the bottlenose dolphin. Future studies will focus on the biology of the ever-growing number of twister ribozymes.

## Conclusion

Overall, the development of CHiTA allowed us to explore the structural diversity of naturally occurring twister ribozymes. We demonstrated that over 1, 900 twisters were extremely tolerant of imperfections in their core and auxiliary helical elements, while tertiary interactions and catalytic residues were sensitive to the type and number of imperfections, allowing some and not others. As such, our structural analysis supports previous findings, but also builds upon them. The structural analysis provided herein better defines the tolerance of twister ribozymes for imperfections, beyond mismatches and mutations. Most importantly, naturally occurring twister ribozymes were found to accommodate an astonishing array of imperfections while maintaining catalytic activity. This analysis can be employed in a recursive fashion to find additional twister candidates. These putative ribozymes could then be tested by CHiTA, to inform ribozyme chemistry and biology, refining the properties that define twister ribozymes. Application of this knowledge will inevitably result in the discovery of additional candidates, like Ttru-1-1 from the bottlenose dolphin. In time, analysis of the genetic context of the twister we tested may help elucidate their biological significance, which remains unclear in many instances. Finally, our analysis can also inform design principles for synthetic applications of ribozymes in the future.

## Acknowledgements

The authors would like to acknowledge the Huck Institutes’ Genomics Core Facility (RRID:SCR_023645). We also thank Kobie Kirven, Reuben Kern, and Dr. Jacob Sieg for their advice on computational and statistical analysis and Dr. Michael Axtell and Dr. Mark Hedglin for helpful comments on the manuscript.

## Funding

This work was supported by the National Institutes of Health [R35-GM127064 to PCB] and a HITS seed grant provided by Huck Institutes of Life Sciences at Penn State.

Will be made available upon peer review of the manuscript.

